# Sem1/DSS1 accelerates ATP-dependent substrate unfolding by the proteasome through a conformation-dependent intercomplex contact

**DOI:** 10.1101/2022.06.27.497739

**Authors:** Randi G. Reed, Gabriel W. Jobin, Robert J. Tomko

## Abstract

The 26S proteasome is an ∼70 subunit ATP-dependent chambered protease that destroys proteins via multiple highly coordinated processing steps. The smallest and only intrinsically disordered proteasome subunit, Sem1 (DSS1 in metazoans), is critical for efficient substrate degradation despite lacking obvious enzymatic activities and being located far away from the proteasome’s catalytic centers. Dissecting its role in proteolysis using cell-based approaches has been challenging because Sem1 also controls proteasome function indirectly via its role in proteasome biogenesis. To circumvent this challenge, we reconstituted Sem1-deficient proteasomes *in vitro* from purified components and systematically dissected its impact on distinct processing steps. Whereas most substrate processing steps are independent of Sem1, ATP-dependent unfolding is stimulated several-fold. Using structure-guided mutagenesis and engineered protein crosslinking, we demonstrate that Sem1 allosterically regulates ATP-dependent substrate unfolding via a distal conformation-dependent intersubunit contact. Together, this work reveals how a small, unstructured subunit comprising < 0.4% the total size of the proteasome can augment substrate processing from afar, and reveals a new allosteric pathway in controlling proteolysis.

## INTRODUCTION

The ubiquitin-proteasome system (UPS) mediates the majority of regulated protein degradation within eukaryotic cells, and defects in the UPS underlie numerous human diseases (reviewed in Chen, et al., 2021). Typically, proteins destined for degradation by the UPS are first modified with a chain of the small protein ubiquitin (Ub). This polyubiquitin (polyUb) chain serves as a targeting signal for delivery to the 26S proteasome (hereafter proteasome). The proteasome is a 2.5 MDa ATP-dependent protease complex that removes the polyUb targeting signal and cleaves the substrate into short peptides. Alterations to the abundance, subunit composition, or function of the proteasome can cause or exacerbate cancers, neurodegenerative disorders, and several autoimmune diseases (Thibaudeau & Smith, 2019). Thus, there is considerable interest in understanding and exploiting proteasome biology for therapeutic benefit.

The proteasome consists of a barrel-shaped 20S core particle capped on one or both ends by the 19S regulatory particle (RP) (**Figure 1A**). The CP comprises four coaxially stacked heptameric rings: two β rings each harboring three different peptidase activities, flanked by two α rings that regulate substrate entry into the proteolytic chamber. The RP can be divided into two subcomplexes, the base and lid (**Figure 1A**). The base consists of a heterohexameric ring of six AAA+ ATPases, Rpt1-Rpt6, and three non-ATPase subunits, Rpn1, Rpn2, and Rpn13. An additional subunit, Rpn10, contacts both the lid and the base and stabilizes their interface. The ATPase subunits are responsible for mechanically unfolding substrates and for opening a gated entryway to the proteolytic chamber of the CP (Eisele, et al., 2018; Rabl, et al., 2008; Smith, et al., 2007; Yedidi, et al., 2017). Rpn1, Rpn10, and Rpn13 act as Ub receptors to recruit substrates (Boughton, Liu, et al., 2021; Husnjak, et al., 2008; Shi, et al., 2016; van Nocker, et al., 1996). The lid consists of nine subunits. Of these, six subunits (Rpn3, Rpn5, Rpn6, Rpn7, Rpn9, and Rpn12) contain α-helical proteasome/cyclosome/initiation complex (PCI) domains, two (Rpn8 and Rpn11) contain Mpr1/Pad1, N-terminal (MPN) domains, and the remaining subunit is the small, intrinsically disordered protein (IDP) Sem1/Rpn15 (DSS1 in mammals). During proteolysis, the Rpn11 deubiquitinating subunit within the lid removes the polyUb targeting signal from the substrate (Worden, et al., 2017), and the ATPase ring of the base mechanically unfolds the substrate and translocates it into the CP for proteolysis.

**Figure 1.**
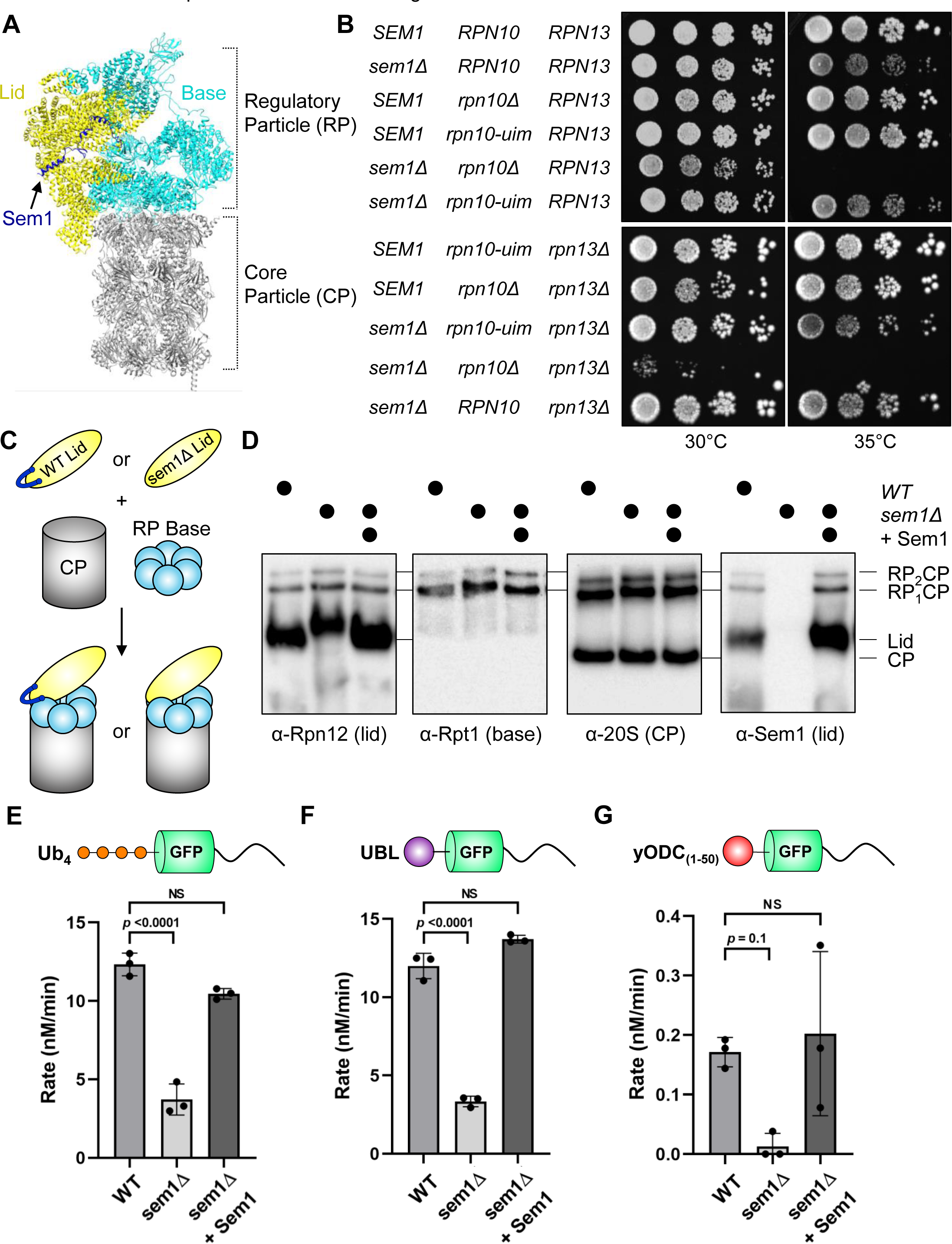
Sem1 lacks genetic interactions with proteasomal substrate-capturing mutations *in vivo* but is essential for efficient substrate degradation *in vitro*. (A) Structure of the 26S proteasome (PDB: 6FVV) (Eisele, et al., 2018) highlighting proteasomal subcomplexes and Sem1. (B) Deletion of *SEM1* fails to yield synthetic growth phenotypes when combined with proteasomal substrate receptor mutations. Equal numbers of cells with the indicated genotypes were incubated on YPD medium for 2 (top) or 3 (bottom) days at the temperatures shown. (C) Schematic of *in vitro* assembly of 26S proteasomes from purified CP, recombinant base, and recombinant lid containing or lacking Sem1. Recombinant Sem1 is added back to sem1Δ proteasomes in some experiments. (D) Proteasomes reconstituted without Sem1 assemble normally and readily incorporate exogenous Sem1. Proteasomes were reconstituted with excess levels of WT lid containing Sem1 (WT) or lid lacking Sem1 (sem1Δ). Recombinant Sem1 was added to sem1Δ proteasomes as shown. After assembly, reconstituted proteasomes were separated by native PAGE and immunoblotted with antibodies against Rpn12, Rpt1, CP, or Sem1. (**E-G**) Degradation of (***E***) Ub_4_-GFP-Tail, (***F***) UBL-GFP-Tail, or (***G***) yODC_(1-50)_-GFP-Tail under multiple-turnover conditions by the indicated reconstituted proteasomes was monitored via loss of GFP fluorescence. For add-back experiments, excess recombinant purified Sem1 was incubated with sem1Δ proteasomes during reconstitution. Shown are the measured degradation rates (N = 3, technical replicates, error bars represent SD). NS, not significant.

Whereas specific contributions to substrate processing have been identified for many proteasome subunits, the functions of several others have remained elusive. This is especially true for Sem1, which was originally identified as a proteasome subunit nearly two decades ago (Funakoshi, et al., 2004; Sone, et al., 2004). However, the function of Sem1 within the proteasome is still poorly defined. At ∼10 kDa, Sem1 is the smallest subunit of the proteasome. Sem1 is largely devoid of secondary and tertiary structure (Paraskevopoulos, et al., 2014), and has several runs of negatively charged amino acids involved in protein-protein interactions. In addition to its role as a proteasome subunit, Sem1 moonlights in several other multisubunit complexes. These include the TREX-2 mRNA-export complex (Faza, et al., 2009) and the Csn12-Thp3 transcription regulating complex (Wilmes, et al., 2008) in budding yeast, and the eIF3 translational initiation complex, the Paf1 and elongator complexes, and the mitotic septin complex in fission yeast (Schenstrøm, et al., 2018). In metazoans, the Sem1 ortholog DSS1 associates with the BRCA2 DNA repair complex (Stefanovie, et al., 2020; Yang, et al., 2002). As Sem1 is seemingly devoid of enzymatic activities, it has generally been thought to serve a scaffolding role in each of these complexes. However, if and how Sem1/DSS1 contributes to the function and regulation of each complex—as well as whether or not it associates transiently or stably—is poorly understood.

The best-understood function of Sem1 within the context of the proteasome is as a facilitator of proteasome lid biogenesis (Tomko & Hochstrasser, 2014). During lid assembly, Sem1 serves an assembly chaperone-like role, in which it tethers subunits Rpn3 and Rpn7 together until their interface can be reinforced by association of additional lid subunits. However, multiple lines of evidence point to assembly-independent roles of Sem1 in proteasomal substrate proteolysis. First, whereas proteasomal assembly chaperones associate transiently with assembling intermediates and dissociate from mature proteasomes (Howell, et al., 2017), Sem1 remains tightly bound to mature proteasomes, seemingly as a stoichiometric component (Bohn, et al., 2013; Funakoshi, et al., 2004; Sone, et al., 2004). Further, although proteasome assembly is compromised in cells lacking Sem1, proteasomes that do form have no overt structural abnormalities (Bohn, et al., 2013). Finally, purified proteasomes lacking Sem1 have a substantial albeit undefined defect in substrate turnover (Funakoshi, et al., 2004; Sone, et al., 2004). Whether, and to what extent, this may stem from altered assembly or more subtle structural defects is not clear.

Recently, Sem1 has been postulated to serve as a Ub receptor for the proteasome based largely on two key observations. First, co-deletion of *SEM1* and the dedicated proteasomal Ub receptor *RPN10* yields a synthetic growth defect in yeast (Funakoshi, et al., 2004; Sone, et al., 2004). Second, purified human DSS1 can bind Ub *in vitro* when the two are mixed (Paraskevopoulos, et al., 2014). However, this Ub receptor function for Sem1/DSS1 has been contested on the basis that: i) co-deletion of *SEM1* and *RPN10* also compromises proteasome assembly and stability that likely accounts for some or all of the synthetic growth defect observed in this double mutant; ii) the regions within Sem1/DSS1 that contact Ub overlap substantially with the sites that contact Rpn3 and Rpn7 within the proteasome, such that simultaneous association of Sem1/DSS1 with the proteasome and with Ub is likely impossible; and iii) no detectable change in substrate binding was observed upon deletion of Sem1 from proteasomes harboring mutations that ablate Ub binding to Rpn10 and Rpn13 (Shi, et al., 2016).

Using cell-based approaches to determine the contribution of Sem1 to proteasomal substrate processing has been challenging because of the confounding roles of Sem1 both in proteasome biogenesis and in the structure and function of other multisubunit complexes. To overcome these limitations, we have reconstituted proteasomes lacking Sem1 *in vitro* from purified components and systematically dissected its role in substrate degradation with a battery of enzymological and biophysical assays. We demonstrate that sem1Δ proteasomes perform virtually all processing steps normally, including capture of substrates, but have greatly reduced substrate unfolding efficiency. This defect results from altered interaction between its binding partner Rpn7 with the proteasomal ATPase ring of the base, and can be fully rescued by provision of ectopic Sem1. Together, our data support a model in which Sem1 allosterically accelerates substrate unfolding through a key lid-base contact, and reveals how a tiny unstructured protein can drive high-efficiency substrate processing by a massive molecular machine ∼250 times its size.

## RESULTS

### Deletion of *SEM1* fails to exacerbate the phenotypes of proteasomal substrate-binding mutants

In budding yeast and likely in other species, Rpn10 is the major Ub receptor for proteasome substrates (Elsasser, et al., 2004; Martinez-Fonts, et al., 2020; Mayor, et al., 2005), and also serves a structural role within the RP (Fu, et al., 2001; Glickman, et al., 1998). Several groups have shown that co-deletion of *RPN10* in *sem1Δ* yeast greatly exacerbates the *sem1Δ* growth defect (Funakoshi, et al., 2004; Krogan, et al., 2004; Sone, et al., 2004; Tomko & Hochstrasser, 2014). The relative contributions of the structural versus the Ub receptor functions of Rpn10 to this phenotype has not been investigated. Toward this goal, we deleted *SEM1* in cells expressing the *rpn10-uim* allele. This allele disrupts the Ub-interacting motif (UIM), thereby allowing Rpn10 to retain its structural role, but preventing its binding to Ub (Elsasser, et al., 2004; Verma, et al., 2004).

Whereas *sem1Δ rpn10Δ* cells were inviable at 35°C, *sem1Δ rpn10-uim* cells grew similarly to *sem1Δ* cells under all conditions tested (**Figure 1B**). To analyze the integrity of these proteasomes, we performed native polyacrylamide gel electrophoresis (native PAGE) on cell lysates from *sem1Δ rpn10Δ* and *sem1Δ rpn10-uim* cells. Consistent with previous observations (Tomko & Hochstrasser, 2014), deletion of *SEM1* alone caused a decrease of doubly capped proteasomes (RP_2_CP) and accumulation of free lid subunit Rpn12, base subcomplex, and CP subcomplex compared to WT cells (**Figure 1 – Figure Supplement 1A**). Deletion of *RPN10* yielded only a modest accumulation of singly RP- capped proteasomes (RP_1_CP) compared to *WT* cells. In contrast, *rpn10-uim* proteasomes assembled similarly to *WT*. Combining *sem1Δ rpn10Δ* resulted in more accumulation of free Rpn12, base, and CP than their individual mutants. However, *sem1Δ rpn10-uim* cells assembled proteasomes similarly to *sem1Δ* cells. Further, we deleted a second Ub receptor, *RPN13*, in cells lacking Sem1 and harboring the Rpn10 Ub-binding mutation. These cells grew similarly to their counterparts containing *SEM1* at 30°C, and similarly to *sem1Δ* cells at 35°C (**Figure 1B**). These data suggest that the synthetic growth defect of *rpn10Δ sem1Δ* cells is due to a structural/assembly issue rather than loss of a redundant substrate capturing function.

Substrates can be recruited directly to proteasomal Ub receptors via a polyUb chain or indirectly via shuttle factors. Shuttle factors use Ub-like (UBL) and Ub-associated (UBA) domains to associate with the proteasome and with substrates, respectively (Bertolaet, et al., 2001; Saeki, et al., 2002; Wilkinson, et al., 2001). In yeast, Rad23 and Dsk2 are the primary proteasomal shuttle factors and preferentially associate with Rpn1 and Rpn13, respectively, on the proteasome (Chen, et al., 2016). As the intrinsic and extrinsic pathways of substrate binding are partly redundant (Boughton, Zhang, et al., 2021; Martinez-Fonts, et al., 2020), we considered that co-deletion of *SEM1* in a *rad23Δ dsk2Δ* background may yield a synthetic defect. However, we found that the triple mutant grew identically to *rad23Δ dsk2Δ* alone (**Figure 1 – Figure Supplement 1B**). Together, these results support previous suggestions (Shi, et al., 2016; Willis, et al., 2020) that, whereas Sem1/DSS1 may bind Ub in isolation, it is unlikely to serve as a Ub receptor in the context of the 26S proteasome.

### Proteasomes reconstituted *in vitro* without Sem1 harbor a substrate degradation defect

The participation of Sem1 in multiple complexes and cellular processes complicates cell-based experiments seeking to understand its function in proteasomal proteolysis. We therefore sought to unequivocally decouple Sem1’s functions in proteolysis from its functions in proteasome assembly and within other complexes. Sem1 is essential for efficient proteasome lid assembly *in vivo* (Tomko & Hochstrasser, 2014); however, we found that under specific conditions we could purify fully assembled recombinant lid lacking Sem1 (hereafter sem1Δ lid) when the remaining eight lid subunits were expressed in *E. coli* (**Figure 1 – Figure Supplement 1C**). We reconstituted proteasomes using purified RP base, CP, and either WT or sem1Δ lid (**Figure 1C**). The integrity of these proteasomes was assessed by native PAGE. Notably, doubly-capped and singly-capped proteasomes formed equally well from either WT or sem1Δ lid. We note that the migration of unincorporated, excess sem1Δ lid subcomplex was slower than that of WT lid (**Figure 1D**), similar to the migration of proteasomes from *sem1Δ* cell extracts (Tomko & Hochstrasser, 2014 and **Figure 1 – Figure Supplement 1A**); this is likely due to a substantial change in the predicted charge state of proteasome lids lacking Sem1 at the pH of electrophoresis (-123.6 for WT versus -109.3 for sem1Δ at pH 8.3). The migration pattern returned to that of WT with the addition of purified Sem1, confirming this hypothesis. No other differences were evident. Combined with previous cryo-EM data on purified *sem1Δ* proteasomes (Bohn, et al., 2013), as well as site-specific crosslinking experiments (see below), we surmise s*em1Δ* proteasomes can assemble efficiently *in vitro* with lid lacking Sem1, presumably because Sem1’s role in lid assembly is “upstream” of lid, base, and CP association (Tomko & Hochstrasser, 2014).

Early studies suggested that proteasomes purified from *sem1Δ* yeast harbor a substantial, but undefined, defect in substrate turnover (Funakoshi, et al., 2004; Sone, et al., 2004). We used the well-characterized model substrate Ub_4_-GFP-Tail (Martinez-Fonts, et al., 2020) to measure substrate degradation. Ub_4_-GFP-Tail contains four linearly fused Ub moieties serving as a proteasomal delivery signal, a circular permutant of superfolder GFP as a reporter for processing, and a disordered region for initiating engagement and unfolding. The loss of GFP fluorescence upon substrate unfolding by the proteasomal ATPases serves as a surrogate reporter for substrate degradation. Consistent with previous studies (Sone, et al., 2004), proteasomes reconstituted without Sem1 displayed a significantly decreased degradation rate compared to WT (**Figure 1E**). Importantly, supplying sem1Δ proteasomes with purified Sem1 restored degradation rates to WT levels, confirming that the absence of Sem1 is solely responsible for the observed degradation defect.

We next tested the ability of sem1Δ proteasomes to degrade two additional substrates that differ from Ub_4_-GFP-Tail only in their proteasomal delivery signal. The first, UBL-GFP-Tail, contains the UBL domain of Rad23 in place of the linear Ub_4_ degron and is preferentially and directly recognized by the T1 substrate receptor site on the proteasome base subunit Rpn1 (Chen, et al., 2016). The second contains the N-terminal 50 amino acids of yeast ornithine decarboxylase (yODC), which binds an as-yet unknown site or sites on the proteasome, but is independent of Ub signaling (Zhang, et al., 2003). Despite containing different degrons, each of these substrates was turned over ≥ 3 times more slowly by sem1Δ proteasomes, and this slowed degradation was completely rescued by addition of recombinant Sem1 (**Figure 1F, G**). Together, these findings confirm that Sem1 enhances substrate degradation by the 26S proteasome, and further indicate that Sem1 enhances substrate degradation independent of the substrate’s delivery signal.

### Reconstituted proteasomes lacking Sem1 display compromised catalytic activity

To better understand the nature of the proteolytic defect in sem1Δ proteasomes, we determined the kinetic parameters of Ub_4_-GFP-Tail degradation by WT and sem1Δ proteasomes. Michaelis-Menten analyses indicated that WT proteasomes degraded Ub_4_- GFP-Tail with a *K*_M_ of 1.5 µM and *k*_cat_ of 0.15 min^-1^, which is similar to what has been previously reported (Martinez-Fonts, et al., 2020; Singh Gautam, et al., 2018). In addition to a small increase in *K*_M_ (∼2-fold), sem1Δ proteasomes displayed an ∼3.8-fold decrease in *k*_cat_, yielding an ∼6.5-fold decrease in catalytic efficiency (*k*_cat_/*K*_M_) compared to WT proteasomes (**Figure 2**). Remarkably, incubating excess purified Sem1 during 26S reconstitution completely restored both *K*_M_ and *k*_cat_ to WT levels (**Figure 2**). Notably, the decrease in *k*_cat_ observed for sem1Δ proteasomes cannot be explained by a simple substrate capture defect because Ub binding is much faster than the subsequent processing steps (Bard, et al., 2019; Lu, et al., 2015). This indicates that Sem1 contributes to one or more downstream catalytic steps in degradation.

**Figure 2.**
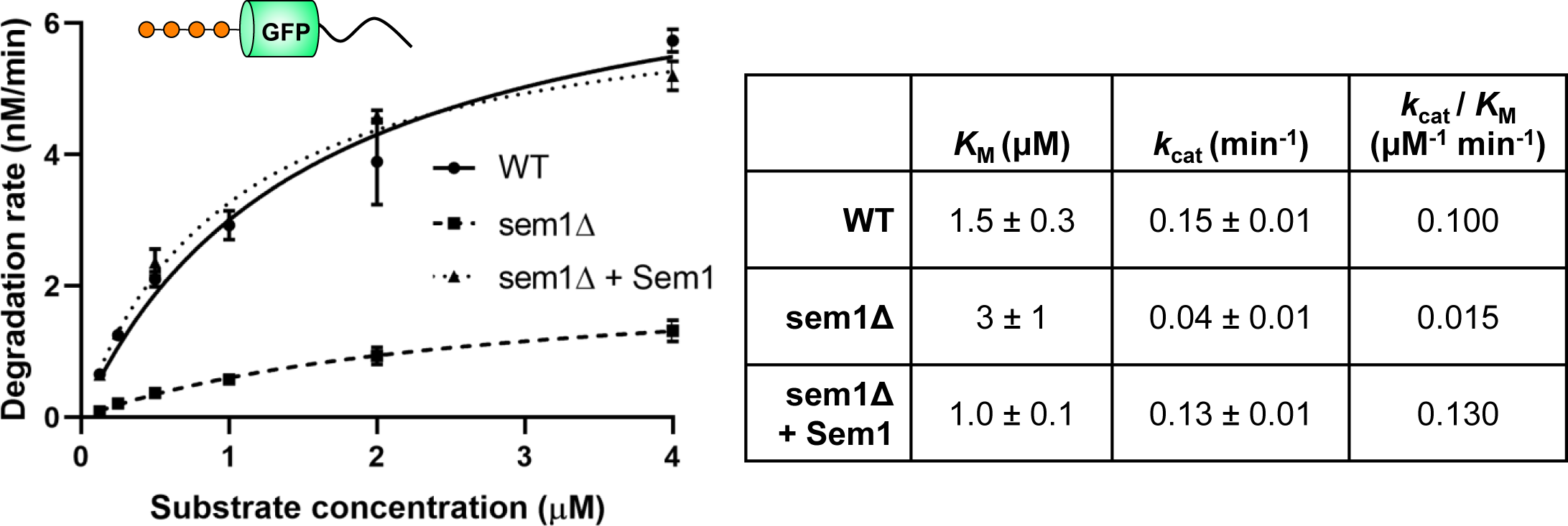
Reconstituted proteasomes lacking Sem1 display compromised catalytic activity. Michaelis-Menten analyses based on initial rates for the turnover of Ub_4_-GFP-Tail by the indicated reconstituted proteasomes. The *K*_M_ and *k*_cat_ values are shown with errors representing SD from the fit (N = 3, technical replicates). The catalytic efficiencies (*k*_cat_ /*K*_M_) are also shown.

### Proteasomes lacking Sem1 display normal substrate engagement, conformational switching, deubiquitination, and peptidase rates

After being captured, a proteasome substrate is first engaged by the proteasomal ATPase pore via an unstructured initiation region (Inobe, et al., 2011; Peth, et al., 2010).

Once engaged, the proteasome undergoes a conformational change that aligns the lid, base, and CP subcomplexes, and poises the deubiquitinase Rpn11 above the ATPase pore (Bard, et al., 2019; Dambacher, et al., 2016; Worden, et al., 2017). In a concerted effort, the ATPases begin mechanically unfolding the substrate to translocate it into the narrow pore of the CP for proteolysis, with polyUb attachments being removed as they are pulled into the active site of Rpn11 by the ATPases (**Figure 3A**).

**Figure 3.**
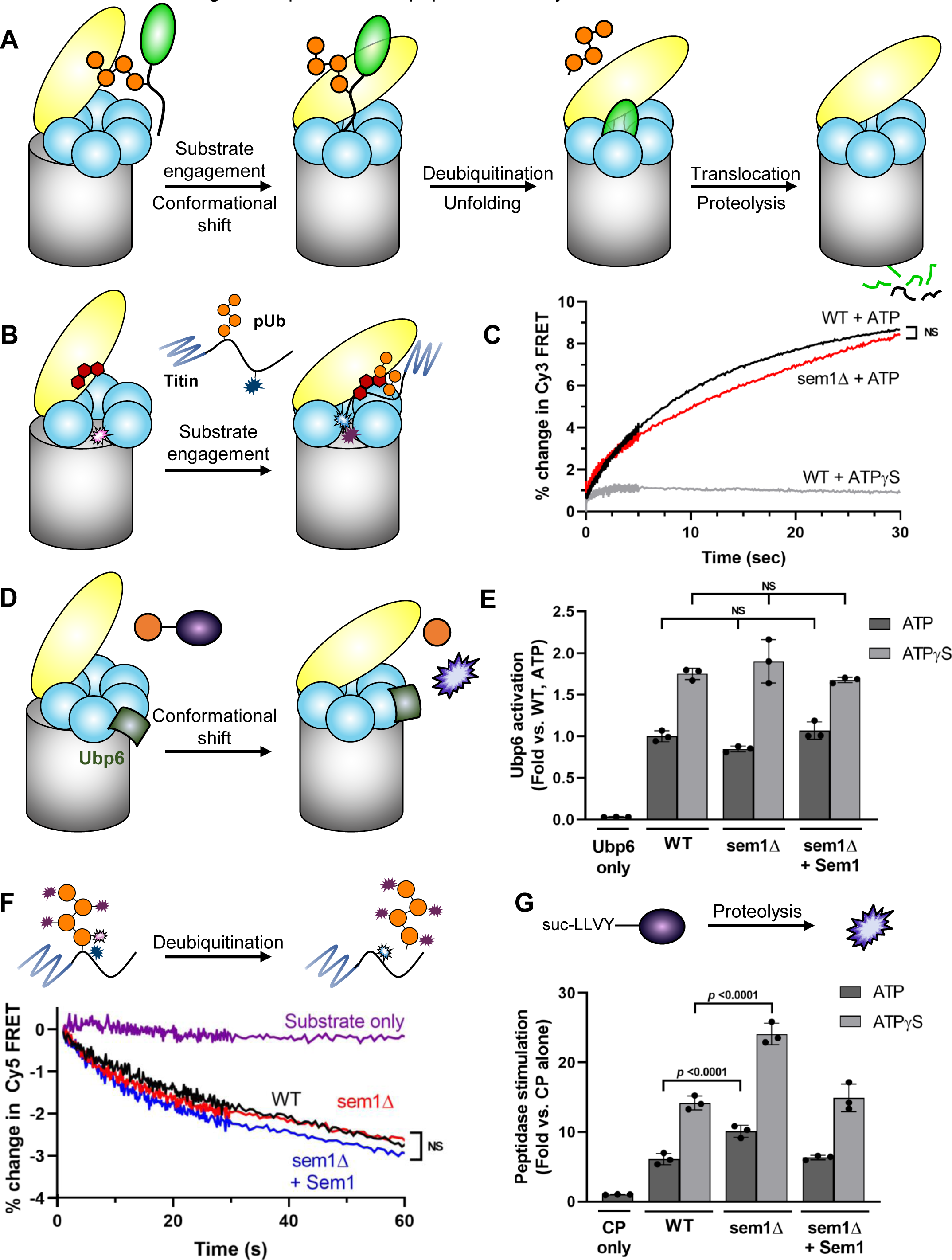
Proteasomes lacking Sem1 have no appreciable defects in substrate engagement, conformational switching, deubiquitination, or peptidase activity. (A) Schematic illustrating the main steps in substrate degradation. Proteasomal subcomplexes are colored as in Figure 1A. A substrate (green oval and black unstructured tail) modified with a polyubiquitin chain (orange circles) illustrates the various processing steps. (B) Schematic depicting a titin-I27^V15P^-Tail substrate labeled with Cy5 (blue star) on its unstructured initiation domain that can undergo FRET with Cy3 (pink star) attached to the central channel of the ATPase ring. In this assay, proteasomes are pretreated with the Rpn11 inhibitor oPA (red) to prevent translocation that would pull the Cy5 molecule on the substrate away from the Cy3 molecule in the ATPase pore, thereby maximizing FRET and enforcing single turnover-like conditions. (C) Representative traces for the change in FRET upon engagement of the substrate tail by WT or sem1Δ proteasomes (N = 3, technical replicates). NS indicates that the difference between rate constants are not significant. (D) Schematic depicting cleavage of Ub-aminomethylcoumarin (AMC) (purple) to Ub and fluorescent AMC after conformation-dependent activation of the extrinsic deubiquitinase Ubp6 (shown in green). NS, not significant. (E) Ubp6 activation was measured via AMC fluorescence in the presence of ATP or ATPγS for the indicated reconstituted proteasomes (N = 3, technical replicates, error bars represent SD). NS, not significant. (F) A titin-I27^V15P^-Tail substrate with Cy5 adjacent to the ubiquitination site is modified with Cy3-labeled ubiquitin. Representative traces for the change in FRET upon deubiquitination of the substrate by the indicated reconstituted proteasomes are shown (N = 3, technical replicates). NS indicates that the differences between rate constants are not significantly significant. (G) The indicated proteasomes were incubated with the fluorogenic peptidase substrate suc-LLVY-AMC. The fold increase in peptidase rates relative to CP alone in the presence of ATP or ATPγS is shown (N = 3, technical replicates, error bars represent SD).

We systematically tested the requirement for Sem1 in each of these steps. First, we employed an established Förster resonance energy transfer (FRET) assay (Bard, et al., 2019) to investigate whether Sem1 affects the engagement of the unstructured substrate tail. In this assay, a donor fluorophore near the central channel of the ATPase pore can excite an acceptor fluorophore positioned in the unstructured tail of a model polyubiquitinated substrate, titin-I27^V15P^-tail (**Figure 3B**), as it is engaged and pulled into the pore. We inhibited deubiquitination by Rpn11 with the Zn^2+^ chelator *ortho*- phenanthroline (Verma, et al., 2004; Wilmes, et al.). Blocking substrate deubiquitination by Rpn11 stalls translocation at the point of ubiquitin attachment, enforcing single-turnover conditions and optimally positioning the acceptor fluorophore for FRET. Tail engagement by WT proteasomes, evident as an increase in FRET, fit well to a one-phase association and occurred with a time constant (τ) of 10.4 s (**Figure 3C, Figure 3 – Figure Supplement 1A-C**). This is longer than previous reports (Bard, et al., 2019), likely due to minor differences in the substrate tail and experimental conditions; however, like previous reports, tail engagement was nearly completely blocked by pre-incubation of WT proteasomes with the slowly hydrolysable ATP analog ATPγS (Bard, et al., 2019). Under the same conditions, sem1Δ proteasomes yielded an increase in FRET with a time constant very similar to that of WT proteasomes (16.4 s). This small difference in tail insertion rate may reflect a decreased overall translocation rate of sem1Δ proteasomes (see below) that slows the threading of the unstructured domain sufficiently for FRET to occur between the donor and acceptor.

We next tested whether Sem1 plays a role in conformational shifting between inactive and active states of the proteasome using a Ubp6 activation assay. The activity of the proteasome-associated deubiquitinase Ubp6 is stimulated upon shifting of the proteasome to an active (also called s3-like) conformational state (Aufderheide, et al., 2015; Bashore, et al., 2015) due to interaction of its USP domain with the proteasomal ATPase ring. The rate of hydrolysis of the fluorogenic substrate Ub-AMC thus provides a surrogate readout of the steady-state conformational distribution of proteasomes between the inactive (also called s1) and activated states (**Figure 3D**). As observed by others (Bashore, et al., 2015; Lee, et al., 2010; Leggett, et al., 2002), pre-incubation of Ubp6 with purified proteasomes and ATP caused a substantial increase in its deubiquitinating activity (**Figure 3E**), and a further increase was observed when ATPγS was provided to bias the distribution toward the active state (Eisele, et al., 2018; Sledz, et al., 2013; Zhu, et al., 2018). Importantly, provision of sem1Δ proteasomes stimulated Ubp6 indistinguishably from WT proteasomes, both in the presence of ATP or ATPγS. Adding purified Sem1 also did not change Ubp6 activation in the presence of either nucleotide. This suggests that Sem1 does not appreciably influence the conformational equilibrium of the proteasome. This observation was further supported by a conformation-selective crosslinking experiment (described below).

Although the Ub_4_-GFP-Tail substrate contains a linear Ub_4_ moiety at its N-terminus, the inclusion of a G76V mutation within the C-terminal diglycine motif of each Ub repeat prevents their cleavage by Rpn11, causing them to be unfolded and degraded with the rest of the substrate. Similarly, both the UBL- and yODC-GFP-Tail substrates lack Ub modifications. As none of these substrates require deubiquitination for degradation, we infer that a deubiquitination defect is not the cause of the slowed degradation by sem1Δ proteasomes. To confirm this, we performed a deubiquitination assay similarly to that previously described (Bard, et al., 2019). Here, we used a titin- I27^V15P^-Tail substrate labeled with an acceptor fluorophore adjacent to its single engineered lysine that, upon polyubiquitination by donor fluorophore-labeled ubiquitin, results in a high FRET signal that can be monitored to measure the kinetics of deubiquitination (**Figure 3F)**. We observed no appreciable difference in the rate of deubiquitination of a polyubiquitinated substrate by WT or sem1Δ proteasomes under single turnover conditions (**Figure 3F, Figure 3 – Supplement 1D-E**), confirming this inference.

We next utilized the fluorogenic peptide proteasome substrate suc-LLVY-AMC to measure whether sem1Δ proteasomes had defects in peptidase activity that may account for the degradation defect. The suc-LLVY-AMC substrate becomes fluorescent when cleaved by the chymotrypsin-like activity of the CP, and conveniently can enter the CP and be cleaved without the need for unfolding or translocation by the ATPase ring. As observed previously (Smith, et al., 2005), provision of ATPγS to WT proteasomes enhanced suc-LLVY-AMC hydrolysis compared to ATP by driving the opening of the proteinaceous gate into the CP (**Figure 3G)**. However, no defect in the peptidase activity of sem1Δ proteasomes relative to WT was observed. In fact, sem1Δ proteasomes displayed a significantly higher basal peptidase rate than WT proteasomes, both when provided ATP or ATPγS (**Figure 3G**). This enhanced peptidase activity could be restored to WT levels by addition of purified Sem1, suggesting that Sem1 negatively regulates CP function. The mechanism by which Sem1 would regulate peptidase activity is not immediately obvious, as Sem1 does not directly contact the CP or the ATPase ring. The simplest explanation is that Sem1 is influencing proteolysis allosterically through interactions of the lid with the CP or ATPase ring (see below). Regardless, we conclude that CP function remains intact in sem1Δ proteasomes.

### Sem1 enhances basal and substrate-stimulated proteasomal ATPase activity

Given that substrate unfolding by the ATPase ring is rate-limiting for WT proteasomes (Bard, et al., 2019) and that most substrate processing steps were relatively unaffected in sem1Δ proteasomes, we tested whether Sem1 influenced proteasomal ATPase activity. We measured a basal ATPase rate for WT proteasomes of ∼34 min^-1^ that was further stimulated 34% to ∼46 min^-1^ upon provision of Ub4-GFP-Tail substrate (**Figure 4A**). These rates are in close agreement with previous reports (Beckwith, et al., 2013; de la Peña, et al., 2018; Snoberger, et al., 2017). In contrast, sem1Δ proteasomes hydrolyzed ATP slower than WT proteasomes, at a rate of ∼26 min^-1^ in the absence of substrate, and were stimulated only 21% to ∼31 min^-1^ upon provision of substrate. Importantly, pre-incubation of sem1Δ proteasomes with recombinant Sem1 prior to measuring ATP hydrolysis restored the observed ATP hydrolysis rates to those observed for WT proteasomes both in the absence and presence of substrate. Together, these observations strongly suggest that Sem1 enhances the ATP hydrolysis rate of both inactive and substrate-bound proteasomes.

**Figure 4.**
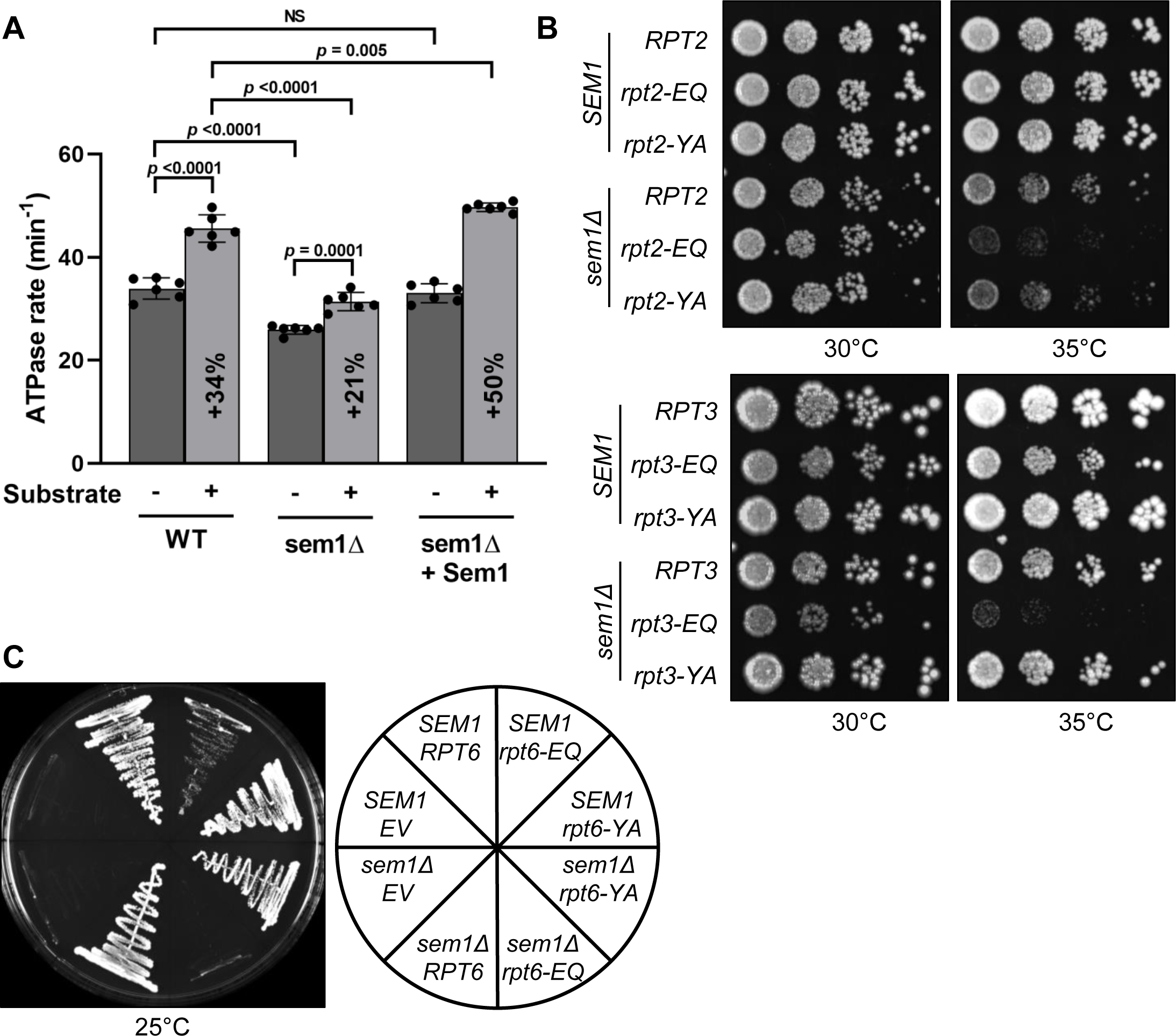
Sem1 positively regulates basal and substrate-stimulated ATP hydrolysis. (A) The ATPase rate of the indicated reconstituted proteasomes was measured in the presence or absence of the substrate Ub_4_-GFP-Tail, with the percent increase upon addition of substrate indicated (N = 6, technical replicates, error bars represent SD). NS, not significant. (B) Equal numbers of *SEM1* or *sem1Δ* cells harboring the indicated WT, ATP hydrolysis mutant (EQ) or pore mutant (YA) *RPT* alleles were spotted in 6-fold serial distributions on YPD and incubated for 2 days (top) or 3 days (bottom) at the indicated temperature. (C) WT or *sem1Δ* yeast cells lacking a chromosomal copy of *RPT6* and sustained with a *URA3-*marked *RPT6* plasmid were transformed with the indicated *RPT6* alleles. Cells were struck on media containing 5-fluoroorotic acid to evict the *URA3-*marked *RPT6* plasmid, and incubated at 25°C for 4 days. The arrangement of strains is shown to the right.

We reasoned that if Sem1 promoted ATP hydrolysis, then combination of *sem1Δ* with ATP hydrolysis mutations in one or more ATPases within the proteasome would yield a synthetic defect. To test this hypothesis, we introduced the well-characterized Walker B EQ substitutions into the Rpt2, Rpt3, and Rpt6 subunits in yeast cells that contained either WT *SEM1* or *sem1Δ* using plasmid shuffling. These mutations have no impact on proteasome assembly (Eisele, et al., 2018), but impair ATP hydrolysis by the subunit bearing the mutation (Beckwith, et al., 2013). Although equivalent Walker B mutations are lethal when introduced into *RPT1*, *RPT4*, or *RPT5*, Sem1 is conveniently most proximal to Rpt2, Rpt3, and Rpt6 and thus most likely to act through one or more of these three adjacent ATPases.

Consistent with a role for Sem1 in accelerating ATP hydrolysis within the proteasome, we found that combining *rpt2-EQ* or *rpt3-EQ* with *sem1Δ* resulted in a synthetic sick phenotype at elevated temperatures (**Figure 4B**). This effect was specific for the Walker B ATP hydrolysis mutants, because combination of *sem1Δ* with *rpt2-YA* or *rpt3-YA* mutants that impair substrate gripping by that particular ATPase (Beckwith, et al., 2013; Erales, et al., 2012) yielded phenotypes no worse than *sem1Δ* alone (**Figure 4B**). This lack of genetic interaction between *sem1Δ* and ATPase pore mutants is also consistent with the lack of a substrate tail insertion defect as described above (**Figure 3B**). A more potent defect was observed for the *sem1Δ rpt6-EQ* mutant, which was synthetic lethal even under ideal growth conditions (**Figure 4C**). This synthetic defect was again specific for the *rpt6-EQ* mutation, as combination of *sem1Δ* with the pore mutant *rpt6-YA* was no worse than *sem1Δ* alone. In sum, these *in vitro* and cell-based experiments indicate an important role for Sem1 in accelerating ATP hydrolysis by the proteasome, and point toward a potential substrate unfolding and/or translocation defect in sem1Δ proteasomes as the source of the degradation defect.

### Sem1 is required for efficient substrate unfolding

To directly assess Sem1’s role in unfolding, we developed a single-turnover unfolding assay inspired by a previous approach utilized to measure Rpn11 activity (Worden, et al., 2017) (**Figure 5A**). In this assay, proteasomes are pre-incubated with the Zn^2+^ chelator *o*PA to inactivate Rpn11. A GFP-Tail substrate harboring a polyUb chain attached to a single lysine between the GFP domain and the unstructured initiation region (**Figure 5 – Figure Supplement 1A**) is then mixed with the *o*PA-treated proteasomes. The substrate is translocated through the ATPase ring until the polyUb branch point reaches the pore, stalling substrate translocation immediately prior to the GFP domain. Rpn11 can then be reactivated by addition of excess Zn^2+^ to permit deubiquitination and re-initiation of unfolding and translocation. As the deubiquitination step is much faster than unfolding (Bard, et al., 2019), the rate of fluorescence loss upon unfolding of the GFP domain directly reflects the unfolding rate. In the absence of Zn^2+^ addition, no decrease in GFP fluorescence was observed (**Figure 5B**), indicating that *o*PA efficiently stalled substrate translocation prior to unfolding of the GFP domain. Upon addition of excess Zn^2+^, stalled WT proteasomes unfolded GFP-tail at a similar rate to proteasomes that were never stalled (**Figure 5B**). In contrast, sem1Δ proteasomes displayed an ∼50% decrease in unfolding rate that was completely rescued by addition of purified Sem1 (**Figure 5B**).

**Figure 5.**
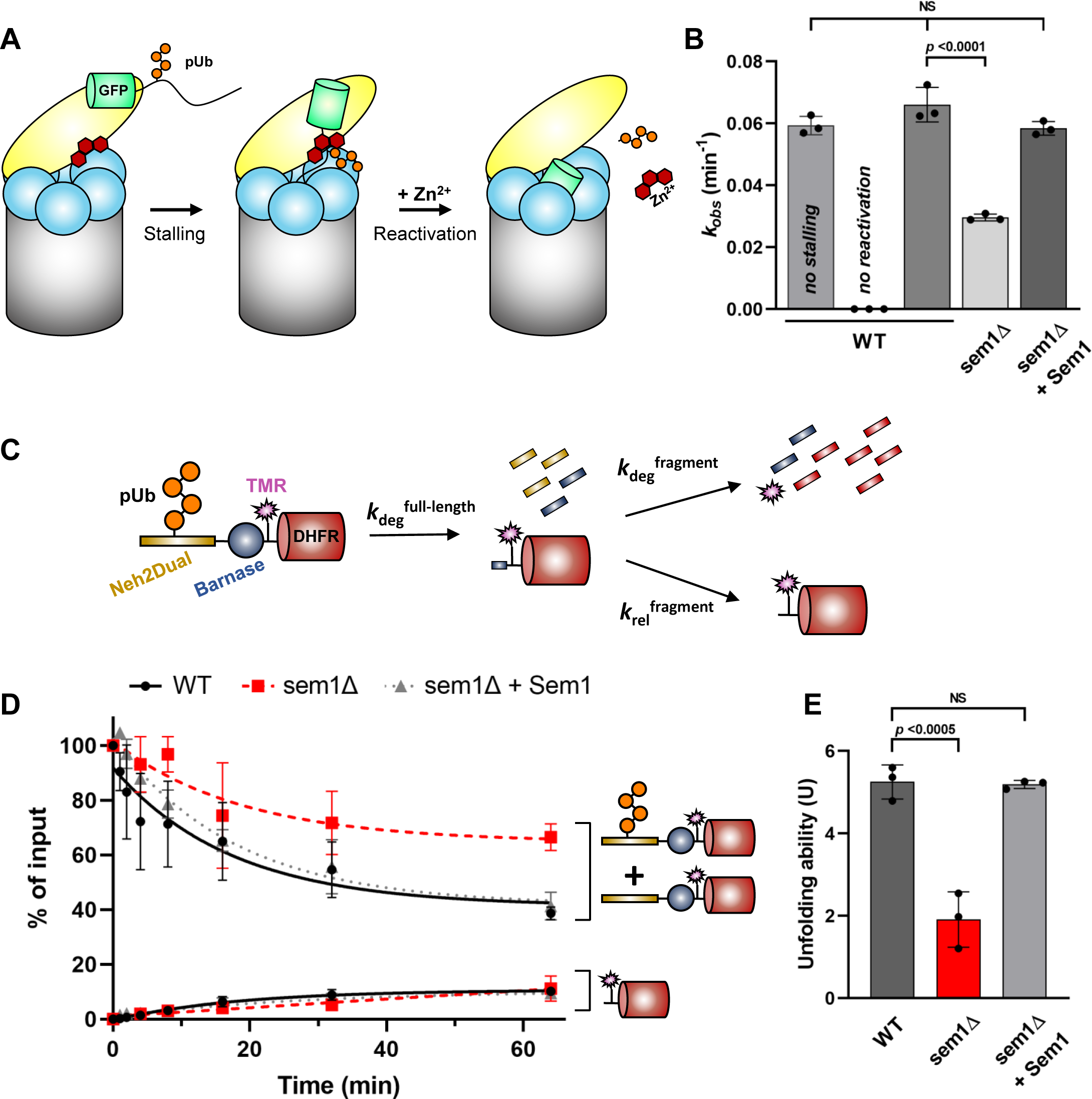
Sem1 is required for efficient substrate unfolding. (A) Schematic illustrating the Rpn11 reactivation assay to measure substrate unfolding. Proteasomes are pre-incubated with the Zn^2+^ chelator *o*PA (red) to inactivate Rpn11 before mixing with a GFP substrate ubiquitinated on its unstructured tail. The polyUb- induced fork in the polypeptide backbone stalls the translocation of substrate immediately before the tightly folded GFP domain is encountered. Addition of excess Zn^2+^ reactivates Rpn11, permitting rapid deubiquitination immediately prior to the comparatively slow unfolding of substrates. (B) The observed unfolding rate constant (*k*_obs_) was determined for the indicated proteasomes by fitting the loss of GFP fluorescence to a one-phase exponential decay (N = 3, technical replicates, error bars represent SD). NS, not significant. (C) Schematic of a model substrate consisting of an N-terminal Neh2Dual domain harboring the sole lysine for polyUb chain attachment, a barnase domain, a site for fluorophore labeling (TAMRA), and a C-terminal DHFR domain. When delivered to the proteasome, the polyUb chain is removed and the substrate is unfolded through the barnase domain at a rate constant *k*deg^full-length^. The tightly folded DHFR domain is either released (*k*rel^fragment^) or processed (*k*deg^fragment^). (D) The amount of remaining full-length substrate (+/- polyUb) or DHFR-containing fragment as a percent of the total input after incubation with the indicated proteasomes was plotted and fit to a one-phase exponential decay or a one-phase exponential association (Figure 5 **– Figure Supplement C-F**) (N = 3, technical replicates, error bars represent SD). (E) The unfolding ability (U) of the proteasomes from (*D*) was determined from the ratio of the rate constants *k*deg^fragment^ and *k*rel^fragment^ (error bars represent SD). NS, not significant.

To confirm this observed unfolding defect in sem1Δ proteasomes, we performed an unfolding ability assay to compare the efficiency with which proteasomes are challenged with a substrate comprising several protein domains that are progressively more difficult to unfold (Cundiff, et al., 2019). This assay utilizes a model substrate consisting of an unstructured N-terminal Neh2Dual initiation region that doubles as a site for polyUb attachment on a single lysine residue, followed by a weakly folded barnase domain, a single fluorophore modification site, and a tightly folded dihydrofolate reductase (DHFR) domain **(Figure 5C, Figure 5 – Figure Supplement 1B**). This DHFR domain can be further stabilized by the addition of NADPH or methotrexate. Proteasomes engage the Neh2 domain, remove the polyUb chain, and unfold and degrade the barnase domain (*k*deg^full-length^). Once the remaining DHFR domain is encountered, the proteasome either unfolds and degrades it (*k*deg^fragment^), or the substrate resists unfolding and is released (*k*rel^fragment^). A lack of a polyUb chain or unstructured initiation region within the DHFR-containing fragment prevents it from being reengaged. The ratio of remaining full-length protein (or deubiquitinated protein) to accumulated DHFR-containing protein fragments is considered the proteasomal unfolding ability (U). As expected, WT proteasomes pre-treated with chemical inhibitors bortezomib, MG132, and *o*PA failed to degrade the full-length substrate (**Figure 5 – Figure Supplement 1F**) whereas untreated WT proteasomes exhibited an U of 5.3 ± 0.2 (**Figure 5D, E, Figure 5 – Figure Supplement 1C**). However, sem1Δ proteasomes displayed a decreased U of 1.9 ± 0.4, which is restored to WT levels (U = 5.2 ± 0.1) upon pre-incubation with recombinant Sem1 (**Figure 5D, E, Figure 5 – Figure Supplement 1D, E**). Taken together with our other observations, we conclude that Sem1 contributes to substrate degradation primarily by enhancing ATP-dependent substrate unfolding.

### Sem1 enhances substrate unfolding via allosteric communication between the lid and base

Sem1 does not directly contact the ATPase ring; rather, the most direct allosteric route between Sem1 and the ATPase ring would be through its interaction with the lid subunit Rpn7 (**Figure 6 – Figure Supplement 1A**). Rpn7 differentially interacts with the ATPase ring in the inactive/s1 state and the activated/s3-like states (Matyskiela, et al., 2013), and close inspection of available high-resolution structures revealed a single site of contact between Rpn7 and Rpt6 in the s3-like states (s2-s6) versus multiple sites in the s1 state. Rpn7 appears to utilize more extensive salt bridging and hydrophobic interactions in the s3-like states versus s1 (**Figure 6A, Figure 7 – Figure Supplement 2A**). We hypothesized that this contact is regulated by Sem1 to drive substrate unfolding. To test this, we introduced mutations at this interface either on Rpn7 (rpn7mut; W204A, E205R, T237A) or on Rpt6 (rpt6mut; R333E, N336A, I375A, H376A) (**Figure 6 – Figure Supplement 1B**). We were unable to obtain the rpt6mut base; we hypothesize this may be due to inadvertent disruption of interaction with the base assembly chaperone Rpn14 (Ehlinger, et al., 2013; Kim, et al., 2010), which is required for base assembly when expressed recombinantly in *E. coli* (data not shown). We moved forward with the rpn7mut lid and assayed Ub4-GFP-Tail degradation by proteasomes before or after addition of ectopic Sem1. The Rpn7 mutations had no impact on the assembly or composition of the lid, and fully assembled into 26S proteasomes *in vitro* (**Figure 6 – Figure Supplement 1C**). When incubated with Ub_4_-GFP-Tail, sem1Δ rpn7mut proteasomes turned over substrate similarly to sem1Δ proteasomes alone; however, addition of recombinant Sem1 failed to restore degradation to WT levels as for sem1Δ proteasomes with WT Rpn7 (**Figure 6B**). We envision that the partial rescue with rpn7mut + Sem1 would have been further suppressed by mutations in Rpt6 at this interface. Regardless, the data support a model in which Sem1 stimulates ATP-dependent substrate unfolding by modulating the Rpn7-Rpt6 contact.

**Figure 6.**
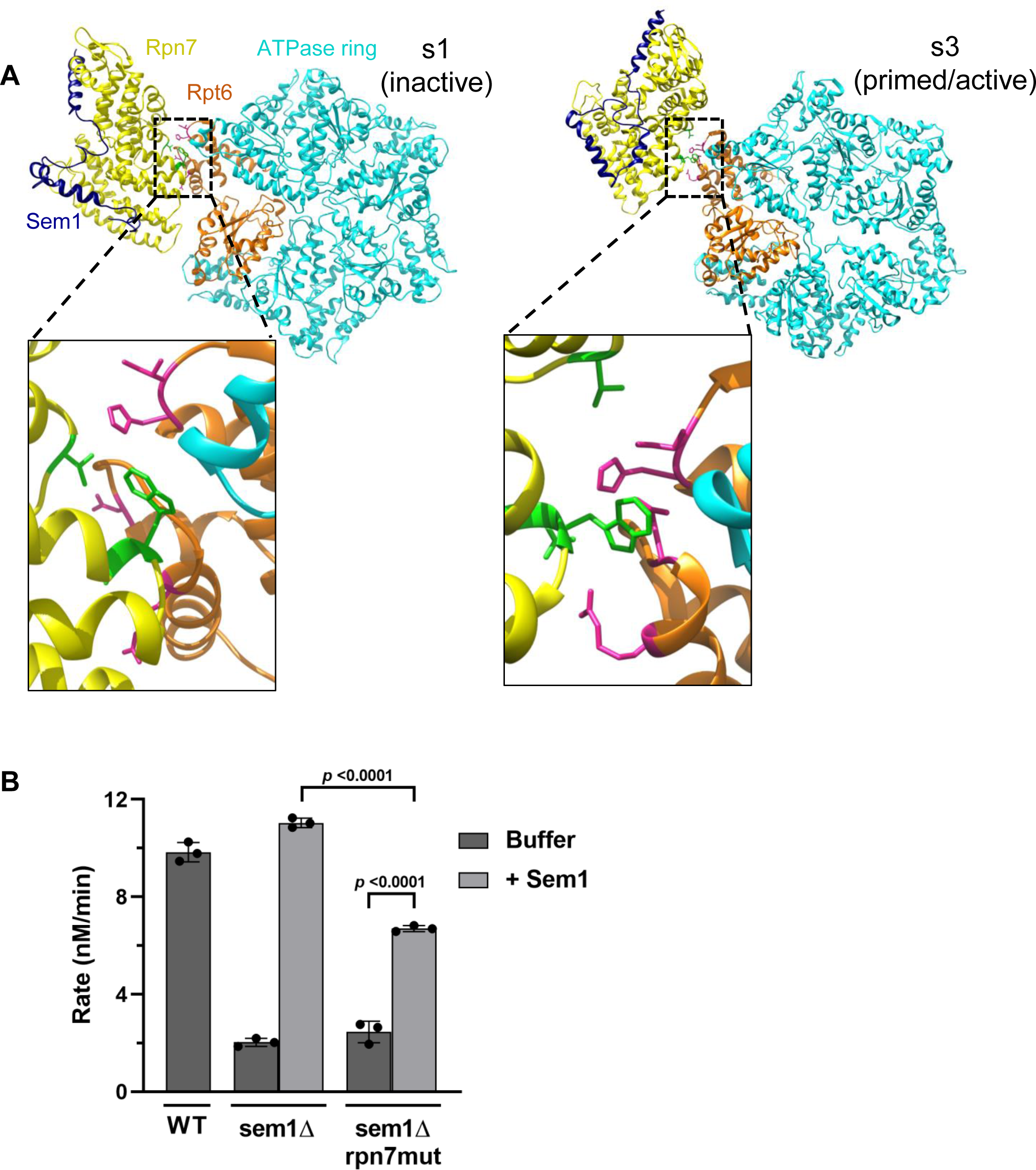
Sem1 allosterically regulates the ATPase activity of the proteasome through Rpn7. (A) Rpn7 forms a single, well-defined contact with Rpt6 in the s3 (primed/active) state (PDB ID: 6FVV), via interactions of Rpn7 residues W204, E205, and T237 (green) with the side chains of Rpt6 amino acids R333, N336, I375, and H376 (pink). In contrast, these Rpt6 side chains are rotated toward a solvent-exposed position in the s1(inactive) state (PDB ID: 6FVT) (Eisele, et al., 2018), with Rpn7 residues instead interacting with the polypeptide backbone. In addition, Rpn7 makes substantial contact with Rpt2 in the s1 state that is completely absent in the s3 state. The coiled-coil and OB fold domains of the ATPase ring are omitted for clarity. (B) Degradation assays using the Ub_4_-GFP-Tail substrate were carried out with the indicated proteasomes reconstituted with or without purified Sem1 (N = 3, technical replicates, error bars represent SD). (C) Proteasomes were reconstituted with excess levels of WT lid containing Sem1 (WT), lid lacking Sem1 (sem1Δ), or rpn7mut lid lacking Sem1 (sem1Δ, rpn7mut). Recombinant Sem1 was added where indicated. After assembly, reconstituted proteasomes were separated by native PAGE and immunoblotted with antibodies against Rpn12, Rpt1, CP, or Sem1.

**Figure 7.**
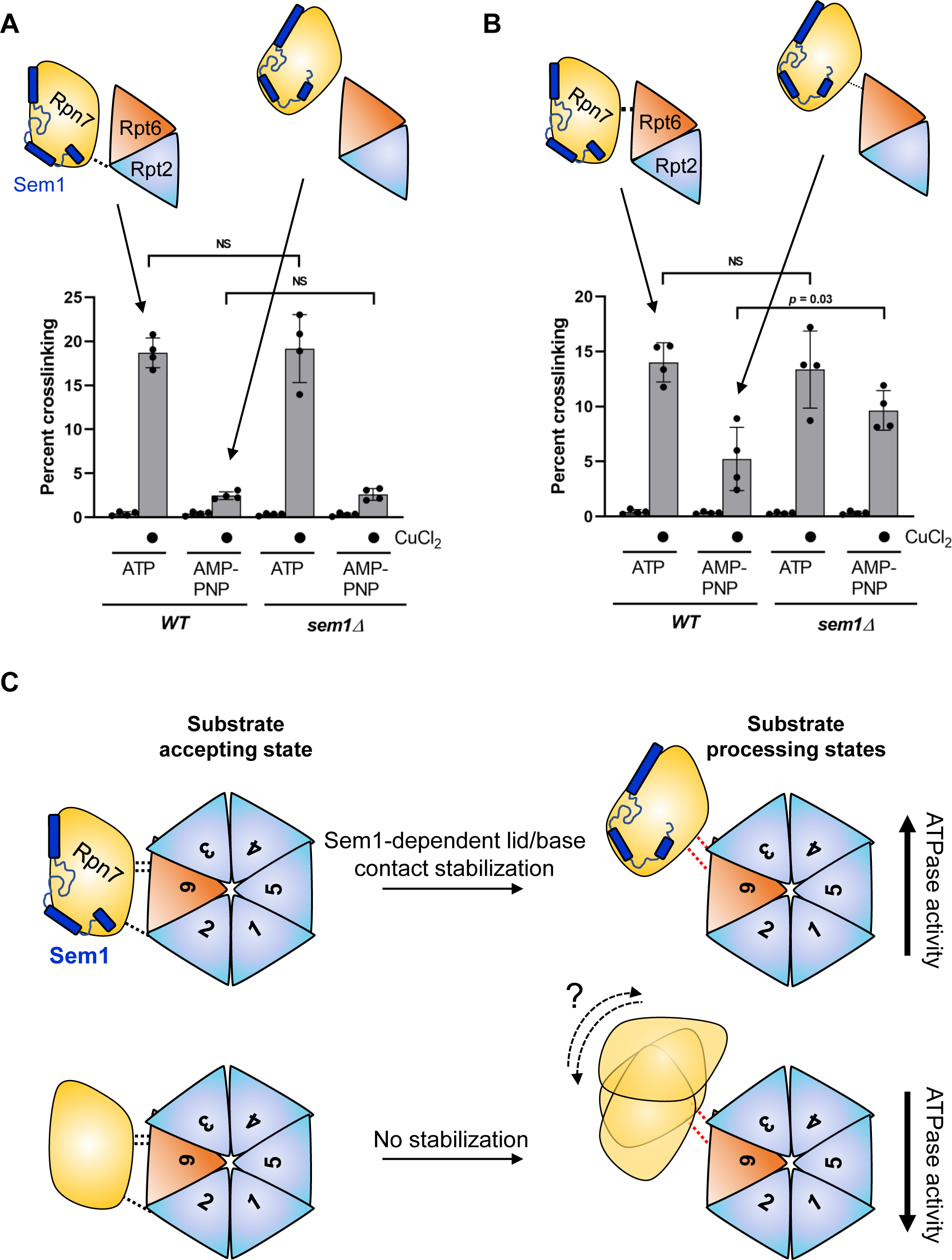
Sem1 stabilizes a lid-base contact critical for ATPase activity and proposed model. (A) Quantitation of crosslinking in WCEs from *WT* and *sem1Δ* cells in the presence of ATP or AMP-PNP in the Rpn7/Rpt2 reporter pair (Figure 7 **– Figure Supplement 1A**). An illustration of the relative positions of Rpn7, Rpt6, and Rpt2 in the s1 state (ATP samples) and the s3-like states (AMP-PNP samples) is shown, with the position of an engineered disulfide crosslink that can be formed in the s1 state shown as a dotted line. (N = 4, technical replicates, error bars represent SD.) NS, not significant. (B) Quantitation of crosslinking in WCEs from *WT* and *sem1Δ* cells in the presence of ATP or AMP-PNP as above, but with the Rpn7-Rpt6 reporter pair (Figure 7 **– Supplement 1C**). The thickness of the dotted line in illustration indicates the formation efficiency of the Rpn7-Rpt6 crosslink in the s1-like (ATP samples) and s3-like (AMP-PNP) states, respectively, when Sem1 is present. (N = 4, technical replicates, error bars represent SD.) NS, not significant. (C) Model for Sem1-dependent control of Rpn7-base interaction. The large- and small- scale conformational changes from the substrate accepting state (s1) into the substrate processing states (s3-like) yields high ATPase activity and efficient substrate degradation. In the absence of Sem1, the small-scale conformational change in the Rpn7-Rpt6 contact is disrupted, either by enabling more dynamic movement of Rpn7, or by failing to fully reposition Rpn7 relative to Rpt6. Disruption of normal Rpn7-Rpt6 interaction in turns leads to reduced ATPase activity and loss of substrate unfolding efficiency.

### Sem1 enhances Rpn7-Rpt6 interaction in the activated state of the proteasome

Considering that: i) Sem1 appears to influence substrate degradation through the Rpn7-Rpt6 contact; and ii) that the Rpn7-Rpt6 contact is remodeled upon proteasome activation (Matyskiela, et al., 2013), we next investigated whether the Rpn7-Rpt6 interface was altered by Sem1, either in the inactive/s1 or activated/s3-like states of the proteasome. We first analyzed the impact of Sem1 on conformational distribution of proteasomes using our previously established conformation-specific crosslinking system that measures the loss of Rpn7 contact with a second ATPase subunit, Rpt2, upon proteasome activation (Eisele, et al., 2018). In this system, cysteine substitutions are made in Rpn7 and Rpt2 such that a disulfide bond can be formed between them in the presence of a mild oxidant such as Cu^2+^. The positions of the substitutions are such that they are close enough to disulfide bond only in the s1 state, but far too distant for disulfide formation in the activated/s3-like states. The abundance of this crosslink thus reports on the proportion of proteasomes in the inactive versus activated states.

The introduction of these cysteine substitutions into *sem1Δ* cells had no apparent effect on overall proteasome structure or cell growth under stress (**Figure 7 – Figure Supplement 1A, B**). Yeast cell lysates were treated with the mild oxidant Cu^2+^, and crosslinks were visualized by non-reducing SDS-PAGE. Consistent with the results of the Ubp6-based conformational assay above, there was no significant difference in the conformational landscape between active and inactive proteasomes using this reporter (**Figure 7A, Figure 7 – Figure Supplement 1C**). Further, the efficiency (abundance) of the Rpn7-Rpt2 crosslink was unaffected by Sem1. Given that disulfide bond formation has a tolerance of only a few Angstroms and is sensitive to the relative orientation of the two sulfhydryl groups, this strongly suggests that the Rpn7-Rpt2 interface remains intact and is remodeled normally during catalysis in the absence of Sem1.

We next engineered a second disulfide crosslinking pair that reports on interaction between Rpn7 and Rpt6 (**Figure 7B, Figure 7 – Figure Supplement 2A**). Again, these substitutions were well tolerated (**Figure 7 – Figure Supplement 2B, C**). As with the Rpn7-Rpt2 reporter pair, crosslinking in the presence of ATP was comparable between *WT* and *sem1Δ* cells. This suggests that, at least in inactive/s1-like proteasomes, no gross defect in the Rpn7-Rpt6 contact point exists. However, when extracts were first incubated with AMP-PNP to drive the proteasomal conformation toward the activated/s3- like states, *sem1Δ* extracts displayed only a minimal decrease in crosslinking compared to ATP, whereas *WT* extracts demonstrated an ∼60% decrease (**Figure 7B, Figure 7 – Figure Supplement 2C**). This observation suggests either: i) a failure to completely remodel the Rpn7-Rpt6 contact during conversion of proteasomes from the inactive/s1 state to the activated/s3-like states; or ii) that the two contact points are more dynamic in the absence of Sem1, resulting in more opportunities to form a disulfide bond during Cu^2+^ treatment. In either case, the interaction between Rpn7 and Rpt6 is altered by Sem1, and this effect is selective for the activated state(s) associated with substrate unfolding and proteolysis. Taken together, these data support a mechanism in which Sem1 allosterically enhances ATP-dependent substrate unfolding through regulation of the Rpn7-ATPase ring interaction.

## DISCUSSION

Since its discovery nearly two decades ago, the role of Sem1 within the proteasome and other macromolecular complexes has remained poorly understood. By leveraging an *in vitro* reconstitution system, we have performed the first detailed mechanistic study of Sem1’s function in proteasomal substrate degradation. We show that Sem1 enhances the efficiency of proteasomal substrate unfolding allosterically through an interaction of the lid subunit Rpn7 with Rpt6 (**Figure 7C**). This post-assembly role for Sem1 rationalizes the impaired substrate degradation observed for sem1Δ proteasomes (Funakoshi, et al., 2004; Sone, et al., 2004) and its persistence within the proteasome after it completes its role in lid assembly (Tomko & Hochstrasser, 2014).

Sem1 has been shown to stabilize and improve the solubility of Thp1 of the TREX- 2 complex in yeast (Ellisdon, et al., 2012) and BRCA2 in mammals (Li, et al., 2006; Yang, et al., 2002). This is a common characteristic of IDPs and intrinsically disordered regions (IDRs) within proteins (Santner, et al., 2012). Specifically, Sem1’s disordered polypeptide chain provides sufficient flexibility for key amino acids within Sem1 to dock onto different surfaces within multisubunit complexes. The highly charged nature of the two acidic patches within Sem1 may function to stabilize the folding of its binding partner. Indeed, another negatively charged and disordered protein, DAXX, was recently demonstrated to utilize its poly-Asp/Glu regions to promote folding of client proteins (Huang, et al., 2021). In agreement with this possibility, we previously found that recombinant expression in *E. coli* of soluble, folded yeast Rpn3 was greatly enhanced by co-expression of Sem1 (Tomko & Hochstrasser, 2014).

It is less common, however, for IDPs or proteins with IDRs to have well-defined molecular functions within a protein or multiprotein complex. IDPs have been reported to serve as flexible linkers to tether larger functional domains (e.g., RPA70 within the RPA complex (Daughdrill, et al., 2007; Jacobs, et al., 1999)), and to serve spring-like functions, such as the IDRs in titin that help to promote relaxation of muscle cells after overstretching (Watanabe, et al., 2002). Sem1 associates with subunits of several multisubunit complexes using distinct binding orientations (Ellisdon, et al., 2012; Tomko & Hochstrasser, 2014; Yang, et al., 2002), gaining different degrees of structure in the various complexes, and even changing its structure during conformational changes in these complexes (Kragelund, et al., 2016). Sem1 may thus act like a tether, spring, or brace depending on the setting.

Our work reveals a new allosteric pathway of communication between the lid and base that is modulated by Sem1 and is crucial for efficient ATP-dependent substrate unfolding. Our group and others recently discovered a role for another conformation-dependent lid-base contact, between Rpn5 and Rpt3. This Rpn5-Rpt3 contact helps to enact conformational changes that both induce the eviction of the base assembly chaperone Nas6 from nascent proteasomes during biogenesis, and activate the proteasome for substrate degradation (Greene, et al., 2019; Nemec, et al., 2019). Although Sem1 also modulates a lid-base contact—Rpn7 with Rpt6—we did not detect any influence on the distribution of proteasomes between these inactive and activated states. Rather, Sem1 appears to reinforce Rpn7, perhaps applying increased pressure against Rpt6 (or other sites) within the ATPase ring to promote optimal ATP hydrolysis and the resultant unfolding and translocation of substrates. Thus, distinct lid-base contacts within the proteasome appear to mediate diverse molecular functions during assembly and catalysis. Further study will be necessary to understand exactly how the Rpn7-Rpt6 contact contributes to ATPase function during catalysis. However, the finding that deletion of *SEM1* is synthetic lethal with the ATPase-dead *rpt6-EQ* mutation argues that it influences ATP hydrolysis more broadly, rather than solely through Rpt6.

Allosteric control of remote macromolecular interactions is emerging as a common functional theme for Sem1. Its human counterpart, DSS1, has recently been suggested to allosterically regulate the position of the tower domain of BRCA2. Although DSS1 binds ∼50 Å away on BRCA2 and makes no contact with this tower, molecular dynamics simulations suggest that docking of DSS1 at this distal site in BRCA2 induces an extended helix-like conformation more conducive to the tower domain’s hypothesized function of capturing dsDNA during homologous recombination (Alagar & Bahadur, 2020; Siaud, et al., 2011; Yang, et al., 2002). We propose that, by stabilizing Rpn7 at a site distal to its contacts with the ATPase ring, Sem1 similarly serves as an allosteric enhancer of the ATPase ring of the proteasome.

DSS1 protects the genome via its role in the BRCA2 complex; however, in apparent contrast to this role, high DSS1 expression increases cellular tolerance to DNA damage independently of BRCA2. In fact, interaction of BRCA2 with DSS1 is dispensable for DNA repair under some conditions (Mishra, et al., 2022). Similarly, breast cancers expressing high levels of DSS1 have worse prognoses and shorter recurrence-free survival times (Gondo, et al., 2021; Rezano, et al., 2013) despite frequently lacking BRCA2. These seemingly paradoxical observations may instead reflect the role of Sem1/DSS1 in the context of the proteasome. The aneuploidy and deregulated gene expression characteristic of cancer cells often renders them dependent on proteasomal proteolysis to avoid cellular damage or death resulting from production of mutant proteins or proteins out of stoichiometry with their normal binding partners (Chen, et al., 2017; Davoli, et al., 2016; Petrocca, et al., 2013; Thibaudeau & Smith, 2019; Walerych, et al., 2016). Although Sem1 is likely a stoichiometric subunit of the proteasome in yeast cells (Funakoshi, et al., 2004; Sone, et al., 2004), it’s unclear whether DSS1 is associated with most or all proteasomes in healthy or cancerous human cells. Enhanced DSS1 expression in these cancers may help maintain protein homeostasis by increasing the unfolding efficiency of proteasomes, yielding improved survival and proliferation. Such a function may explain, in whole or in part, the aggressive nature and drug resistance of DSS1-overproducing cancers.

It remains to be seen whether Sem1/DSS1 is a druggable target for antineoplastic therapy. Although IDPs are traditionally considered difficult to target with small molecules due to the lack of defined conformations and binding pockets, inhibitors have recently been discovered for several IDPs involved in cancers and neurodegenerative diseases (Santofimia-Castaño, et al., 2020). Given that upwards of ∼50% of proteins in eukaryotic proteomes are predicted to have disordered regions ≥ 40 consecutive amino acids in length (Dunker, et al., 2002), understanding how IDPs and IDRs influence biological processes could potentially illuminate many new targets for therapeutic intervention in human disease.

## METHOD DETAILS

### Yeast strains and media

All yeast strains used in this study are listed in Table I. Yeast strains were generated and handled according to standard protocols (Guthrie & Fink, 1991). All yeast strains were grown in YPD medium at 30°C, or in the case of *RPT* Walker B mutants and their respective controls, at 25°C. To select for plasmids, strains were grown in synthetic dropout medium lacking the appropriate auxotrophic agent. Growth assays were conducted by spotting equal numbers of cells in six-fold serial dilutions in water onto the indicated media.

**Table I:** Yeast strains used in this study.

| **Name** | **Genotype** | **Source** |
| --- | --- | --- |
| RTY1 | *MATa his3-Δ200 leu2-3,112 ura3-52 lys2-801 trp1-1*<br>(alias MHY500) | (Chen, et al., 1993) |
| RTY20 | *MATa his3-Δ200 leu2-3,112 ura3-52 lys2-801 trp1-1*<br>*rpn10Δ::HIS3* (alias MHY960) | (Tomko & Hochstrasser, 2011) |
| RTY21 | *MATα his3-Δ200 leu2-3,112 ura3-52 lys2-801 trp1-1*<br>*rpn10Δ::HIS3 sem1Δ::HIS3* | This study |
| RTY85 | *MATa his3-Δ200 leu2-3,112 ura3-52 lys2-801 trp1-1*<br>*sem1Δ::HIS3* (alias MHY4785) | (Tomko & Hochstrasser, 2011) |
| RTY691 | *MATa his3-Δ200 leu2-3,112 ura3-52 lys2-801 trp1-1*<br>*rpn10-uim:kanMX* | This study |
| RTY975 | *MATa his3-Δ200 leu2-3,112 ura3-52 lys2-801 trp1-1*<br>*rpn10-uim:kanMX sem1Δ::HIS3* | This study |
| RTY1821 | *MATa his3-Δ200 leu2-3,112 ura3-52 lys2-801 trp1-1*<br>*rpt2Δ::hphMX4 [pRS316-RPT2]* | This study |
| RTY1823 | *MATa his3-Δ200 leu2-3,112 ura3-52 lys2-801 trp1-1*<br>*rpt3Δ::hphMX4 [YCplac33-RPT3]* | This study |
| RTY1827 | *MATa his3-Δ200 leu2-3,112 ura3-52 lys2-801 trp1-1*<br>*rpt6Δ::hphMX4 [YCplac33-RPT6]* | This study |
| RTY2091 | *MATa his3-Δ200 leu2-3,112 ura3-52 lys2-801 trp1-1*<br>*rpn7(D123C)-6xGly-V5:kanMX6* | (Eisele, et al., 2018) |
| RTY2099 | *MATa his3-Δ200 leu2-3,112 ura3-52 lys2-801 trp1-1*<br>*rpt2(R407C):natMX4* | (Eisele, et al., 2018) |
| RTY2112 | *MATa his3-Δ200 leu2-3,112 ura3-52 lys2-801 trp1-1*<br>*rpn7(D123C)-6xGly-V5:kanMX6 rpt2(R407C):natMX4* | (Eisele, et al., 2018) |
| RTY2156 | *MATa his3-Δ200 leu2-3,112 ura3-52 lys2-801 trp1-1*<br>*rpn10-uim:kanMX rpn13Δ::HIS3MX6* | This study |
| RTY2529 | *MATa his3-Δ200 leu2-3,112 ura3-52 lys2-801 trp1-1*<br>*rpn7(D123C)-6xGly-V5:kanMX6 rpt2(R407C):natMX4*<br>*sem1Δ::HIS3* | This study |
| RTY2534 | *MATa his3-Δ200 leu2-3,112 ura3-52 lys2-801 trp1-1 PRE1-6xGly-6His:kanMX6 pep4Δ::NAT* | This study |
| RTY2772 | *MATa his3-Δ200 leu2-3,112 ura3-52 lys2-801 trp1-1 rpt6Δ::hphMX4 [YCplac33-RPT6] sem1Δ::HIS3* | This study |
| RTY2857 | *MATa his3-Δ200 leu2-3,112 ura3-52 lys2-801 trp1-1 rpt2Δ::hphMX4 [pRS316-RPT2] sem1Δ::HIS3* | This study |
| RTY2859 | *MATa his3-Δ200 leu2-3,112 ura3-52 lys2-801 trp1-1 rpt3Δ::hphMX4 [YCplac33-RPT3] sem1Δ::HIS3* | This study |
| RTY2980 | *MATa his3-Δ200 leu2-3,112 ura3-52 lys2-801 trp1-1 dsk2Δ::natMX4* | This study |
| RTY2983 | *MATα his3-Δ200 leu2-3,112 ura3-52 lys2-801 trp1-1 rad23Δ::kanMX6* | This study |
| RTY3111 | *MATα his3-Δ200 leu2-3,112 ura3-52 lys2-801 trp1-1 dsk2Δ::natMX4 rad23Δ::kanMX6* | This study |
| RTY3147 | *MATa his3-Δ200 leu2-3,112 ura3-52 lys2-801 trp1-1 dsk2Δ::natMX4 rad23Δ::kanMX6 sem1Δ::HIS3* | This study |
| RTY3630 | *MATa his3-Δ200 leu2-3,112 ura3-52 lys2-801 trp1-1 rpn7Δ::natMX4 [pRS316-RPN7] rpt6Δ::hphMX4 [YCplac33-RPT6] sem1Δ::HIS3* | This study |
| RTY3632 | *MATa his3-Δ200 leu2-3,112 ura3-52 lys2-801 trp1-1 rpn7Δ::natMX4 [pRS316-RPN7] rpt6Δ::hphMX4 [YCplac33-RPT6]* | This study |
| RTY3722 | *MATa his3-Δ200 leu2-3,112 ura3-52 lys2-801 trp1-1 rpn10Δ::hphMX4 rpn13Δ::HIS3MX6* | This study |
| RTY3724 | *MATa his3-Δ200 leu2-3,112 ura3-52 lys2-801 trp1-1 rpn13Δ::HIS3MX6 sem1Δ::HIS3* | This study |
| RTY3726 | *MATa his3-Δ200 leu2-3,112 ura3-52 lys2-801 trp1-1 rpn10-uim:kanMX rpn13Δ::HIS3MX6 sem1Δ::HIS3* | This study |
| RTY3730 | *MATa his3-Δ200 leu2-3,112 ura3-52 lys2-801 trp1-1 rpn10Δ::hphMX4 rpn13Δ::HIS3MX6 sem1Δ::HIS3* | This study |

### Plasmids

Plasmids used in this study are listed in Table II. All plasmids were constructed using standard molecular cloning techniques with TOP10 F’ (Thermo-Fisher) as a host strain. QuikChange (Agilent) was used for site-directed mutagenesis. All open reading frames were sequenced prior to use. Complete plasmid sequences and construction details are available upon request.

**Table II:**
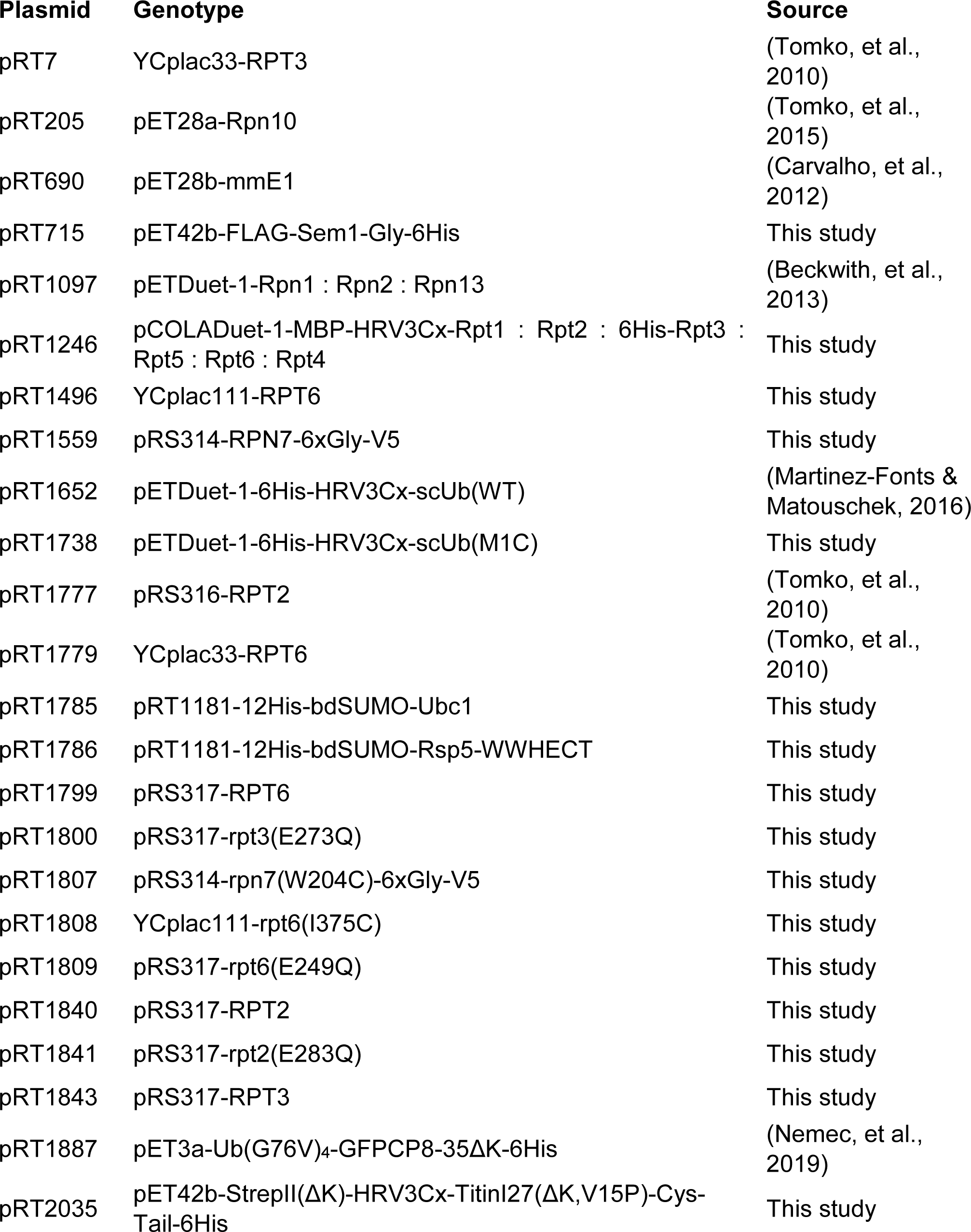
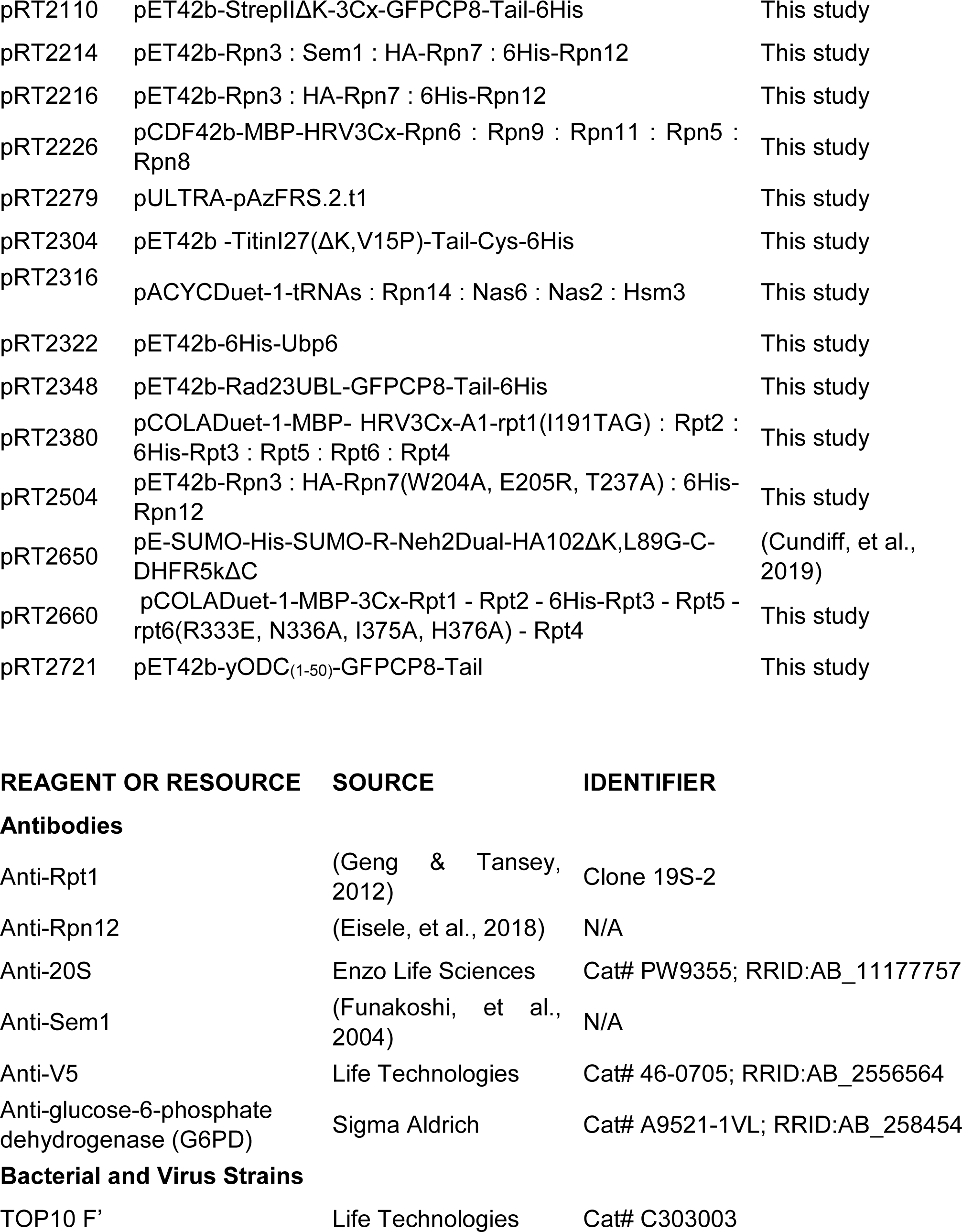
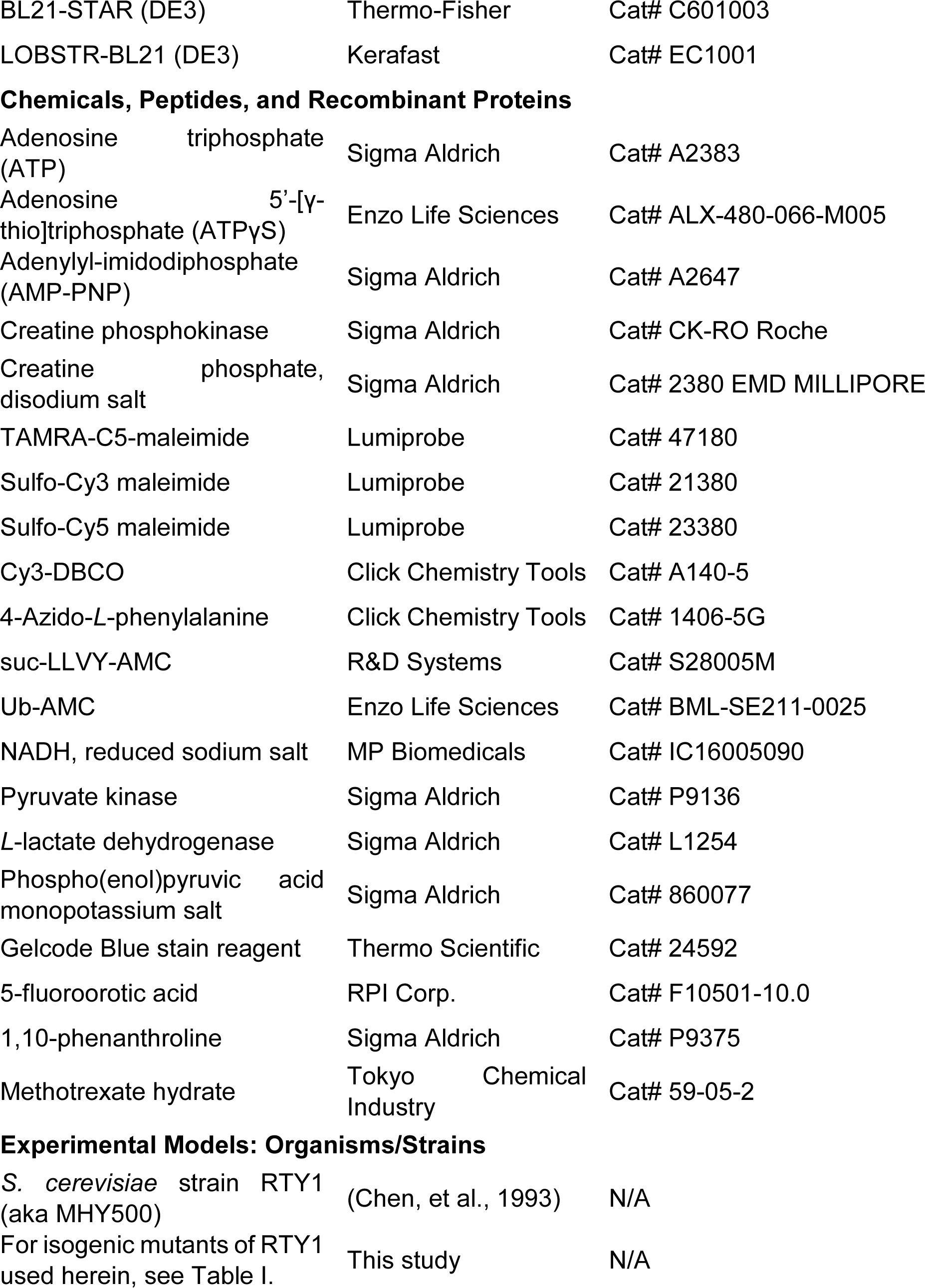
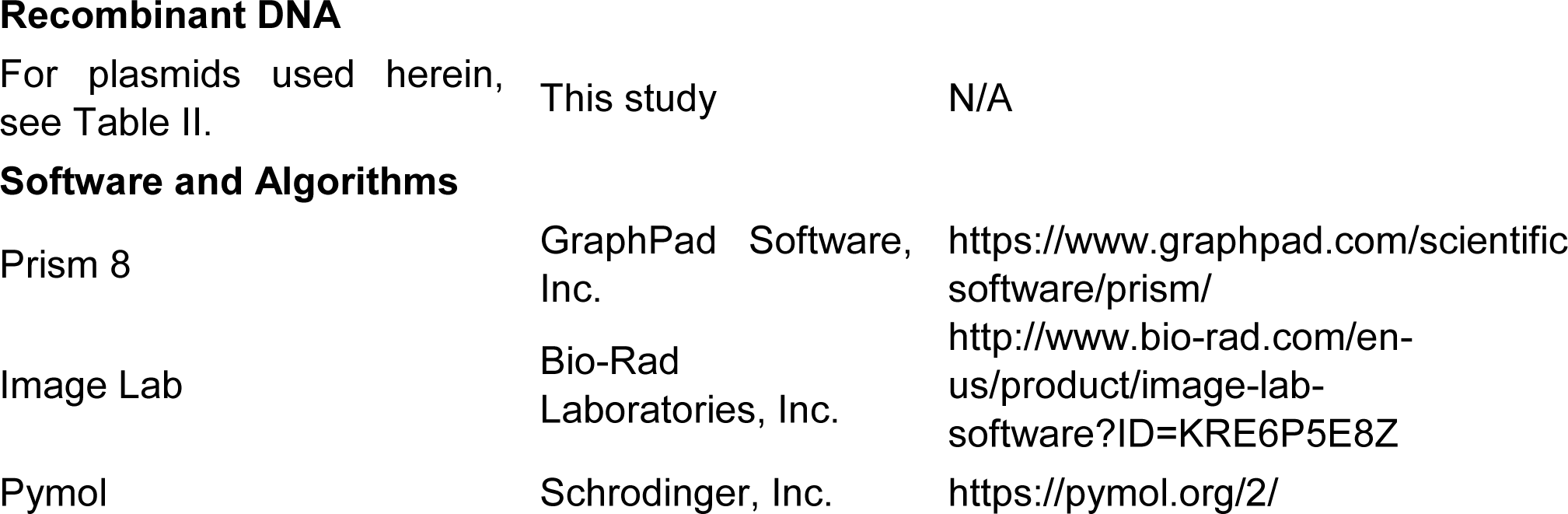
Plasmids used in this study.

### Non-denaturing polyacrylamide gel electrophoresis

For analysis of cell extracts, 60 mg of total protein was separated by 4% non-denaturing PAGE as described previously (Nemec, et al., 2017). Specifically, cells were grown to OD_600_ ∼2.0, harvested by centrifugation at 8,200 x *g* for five minutes at room temperature, followed by washing in 25 mL of ice-cold water. Cells were centrifuged again at 5,000 x *g* for two minutes, 4°C, and the supernatant was decanted. Cell pellets were then frozen in liquid nitrogen and ground into powder using a mortar and pestle. Cell powder was hydrated in one powder volume of extraction buffer (50 mM Tris•Cl, pH 7.5, 5 mM MgCl_2_, 10% glycerol, 1 mM ATP, 0.015% w/v xylene cyanol), and incubated with frequent vortexing on ice for 10 minutes. Cell debris was removed by centrifugation at 21,000 x *g*, 4°C for 10 minutes. Supernatants containing equal amounts of protein (determined by BCA assay) were loaded onto 4% native polyacrylamide gels cast with 0.5 mM ATP and with a 3.5% polyacrylamide stacking gel containing 2.5% sucrose and 0.5 mM ATP. Electrophoresis was conducted at 100 V, 4°C until the dye front escaped.

For analysis of purified components, 20 nM CP, 40 nM base, 40 nM Rpn10, and 200 nM lid were combined in extraction buffer and allowed to assemble at 30°C for 15 minutes prior to native PAGE as above.

### Immunoblot analyses

Proteins separated by native PAGE or SDS-PAGE were transferred to PVDF membranes and immunoblotted with antibodies against Rpn12 (Eisele, et al., 2018) (1:10,000), 20S CP (Enzo Life Sciences Cat# PW8160) (1:1250), Rpt1 19S-2 (Geng & Tansey, 2012) (1:5000), Sem1 (Funakoshi, et al., 2004) (1:2000), V5 (Life Technologies Cat# 46-0705) (1:5000), or glucose-6-phosphate dehydrogenase (G6PD) (Sigma Cat#A9521-1VL) (1:20,000). Blots were imaged using a Bio-Rad ChemiDoc MP using HRP-conjugated secondary antibodies (Cytiva Life Sciences) and ECL reagent.

### Protein Purifications

Most proteins underwent a final gel filtration purification step, and the purest fractions were pooled, concentrated using appropriate centrifugal filters, and snap-frozen as small aliquots in liquid nitrogen for storage at -80°C.

### Purification of CP

CP was purified from yeast strain RTY2534 via ammonium precipitation, Ni-NTA affinity, anion exchange, and gel filtration as previously described (Nemec & Tomko, 2020). Six liters of YPD inoculated with RTY2534 were grown until saturation and were harvested at 5000 x *g*, 25°C for five minutes. The cell pellet was washed once in 250 mL of deionized water and snap-frozen in liquid nitrogen. The frozen cells were then ground into powder using a SPEX 6850 freezer mill and stored at -80°C. The day of purification, the cell powder was thawed in an equal volume of CP buffer (50 mM Tris•Cl, pH 7.5, 50 mM NaCl, 50 mM KCl, 0.5 mM EDTA) supplemented with 0.05% NP-40 and stirred until thawed. The insoluble debris was pelleted at 30,000 x *g*, 4°C for 20 minutes. Ammonium sulfate was slowly added to the supernatant to 40% saturation, and the mixture was stirred for 1 hour at 4°C before being centrifuged again. The supernatant was then slowly adjusted to 70% ammonium sulfate and centrifuged again as before. The pellet was resuspended in CP Ni Binding Buffer (50 mM Tris•Cl, pH 7.5, 500 mM NaCl, 10 mM imidazole), added to Ni-NTA resin, and allowed to incubate for 45 minutes at 4°C. The resin was collected at 1500 x *g*, 4°C for two minutes, and the supernatant was decanted. The resin was then washed with Ni CP Binding Buffer followed by CP Ni Low Salt Wash Buffer (50 mM Tris•Cl, pH 7.5, 50 mM NaCl, 10 mM imidazole). CP Elution Buffer (50 mM HEPES•OH, pH 7.5, 100 mM NaCl, 100 mM KCl, 5% glycerol, 500 mM imidazole) was added to the resin to elute protein. The eluate was loaded onto a Mono Q 5/5 column equilibrated in CP IX-A Buffer (50 mM Tris•Cl, pH 7.5, 5 mM MgCl_2_•6H_2_O), and proteins were eluted by a continuous gradient of CP IX-B Buffer (50 mM Tris•Cl, pH 7.5, 1 M NaCl, 5 mM MgCl_2_•6H_2_O). The purest enzymatically active fractions were pooled and concentrated in a 30,000 Da MWCO Amicon filter, and further purified by gel filtration on a Superose 6 10-30 column equilibrated in CP buffer.

### Purification of recombinant base complexes

Recombinant base was expressed from plasmids pRT2316, pRT1097, and pRT1246 (or pRT2660 for rpt6mut base) co-transformed into bacterial strain BL21-STAR (DE3) following a modified version of a published method (Beckwith, et al., 2013). Specifically, transformants were grown in terrific broth and the appropriate antibiotics at 37°C to an OD_600_ ≈ 0.7, at which point the temperature was reduced to 30°C for 30 minutes. IPTG was then added to 0.5 mM and the temperature was reduced further to 16°C for overnight induction. Cultures were centrifuged at 8200 x *g*, 16°C for 5 minutes. The cell pellet was resuspended in Base Buffer (50 mM HEPES•OH, pH 7.5, 100 mM NaCl, 100 mM KCl, 10 mM MgCl_2_•6H_2_O, 0.5 mM ATP) containing 20 mM imidazole and supplemented with 0.1% NP-40, 1 mg/mL lysozyme, 3.3 U/mL benzonase, and 1 mM PMSF before being frozen at -80°C until purification. The day of purification, cells were thawed and lysed by sonication. The lysate was clarified by centrifugation at 30,000 x *g*, 4°C for 20 minutes, and added to Ni-NTA resin for 45 minutes. After washing with Base Buffer containing 20 mM imidazole, proteins were eluted with Base Buffer containing 250 mM imidazole. The eluate was then incubated with amylose resin for 45 minutes at 4°C, washed twice with Base Buffer supplemented with 1 mM DTT, and eluted with Base Buffer containing 20 mM *D-*maltose. The eluate was concentrated to a small volume (<500 µL) in a 100,000 Da MWCO Amicon filter, and the MBP tag on Rpt1 was cleaved with 1:20 (w/w) HRV-3C protease overnight at 4°C. The following morning, the sample was purified by gel filtration on a Superose 6 10-30 column in Base Buffer supplemented with 500 µM TCEP. The concentration of base was determined by BCA assay using bovine serum albumin (BSA) as a standard.

Recombinant base containing 4-azido-*L*-phenylalanine (AzF) was expressed, purified, and labeled similarly to above with the following modifications (Bard, et al., 2019). BL21-STAR (DE3) was co-transformed with plasmid pRT2279 harboring the AzF tRNA synthetase/tRNA pair, pRT2316, pRT1097, and pRT2380 (harboring the Rpt1 I-191-TAG mutation). Transformants were grown in 6 L 2xYT media until OD_600_ ≈ 1.4, and cells were spun down at 3500 x g for 10 minutes, 30°C. The cells were resuspended in UAA media (24 g yeast extract, 20 g tryptone, 1% glycerol, 17 mM KH_2_PO_4_, 72 mM K_2_HPO_4_ for 1 L) supplemented with 2 mM AzF, and incubated with shaking at 30°C for 30 minutes to allow for uptake of the AzF. Cultures were then induced with 0.5 mM IPTG for 5 hours at 30°C before incubation at 16° overnight. Purification was conducted as above, except with no reducing agent. After Ni-NTA and MBP affinity purifications as above, the eluate was concentrated to a small volume (∼350 µL). Free thiols were blocked by the addition of 150 µM 5,5’-dithio-bis-(2-nitrobenzoic acid) for ten minutes at room temperature. The eluate was returned to 4°C and labeled overnight with 300 µM dibenzocyclooctyne- conjugated Cy3. The reaction as quenched with 1.5 mM sodium azide for 20 minutes on ice. The MBP tag was cleaved with 1:20 (w/w) HRV-3C protease in the presence of DTT for 1 hour at room temperature. The sample was then purified by gel filtration, and the concentration was determined as above.

### Purification of recombinant lid complexes

Fully recombinant lid complex and sem1Δ lid were expressed from pRT2226 and either pRT2214 (WT lid), pRT2216 (sem1Δ lid), or pRT2504 (sem1Δ, rpn7mut lid) as previously described (Nemec, et al., 2019). Lid was expressed in LOBSTR (DE3) co-transformed with pRARE2. Transformants were grown in terrific broth and the appropriate antibiotics at 37°C until OD_600_ ≈ 1.0, at which point the temperature was reduced to 16°C and IPTG was added to 0.5 mM. After overnight induction, cultures were centrifuged at 8200 x *g*, 16°C for 5 minutes, the supernatant was poured off, and the cells were stored at -80°C. The day of purification, the cell pellet was thawed in Low Salt Lid (50 mM HEPES•OH, pH 7.5, 50 mM NaCl, 5% glycerol) supplemented with 0.1% NP-40, 5 mM β-mercaptoethanol, and 1 mM PMSF, and the cells were lysed with an Avestin Emulsiflex C-5. Lysates were clarified via centrifugation at 30,000 x *g*, 4°C for 20 minutes, and the supernatant was incubated with amylose resin for 45 minutes at 4°C. After two washes with Low Salt Lid Buffer supplemented with 5 mM β-ME, proteins were eluted with Low Salt Lid Buffer supplemented with 20 mM *D-*maltose. The eluate was incubated with Ni-NTA resin for 45 minutes at 4°C. After two washes with Low Salt Lid Buffer supplemented with 5 mM β-ME and 10 mM imidazole, proteins were eluted by increasing imidazole concentration to 250 mM. The eluate was concentrated to <500 µL in a 100,000 Da MWCO Amicon filter, and the MBP tag on Rpn6 was cleaved by incubating with 1:20 (w/w) HRV-3C protease overnight at 4°C. The following morning, the sample was purified by gel filtration on a Superose 6 10-30 column in Low Salt Lid Buffer.

### Purification of Rpn10

Rpn10 was expressed from pRT205 as an N-terminal 6His fusion in bacterial strain LOBSTR (DE3) co-transformed with pRARE2 as previously described (Nemec, et al., 2019). Transformants were grown in LB and the appropriate antibiotics at 37°C, 250 rpm shaking until OD_600_ ≈ 0.6, at which point the temperature was reduced to 30°C and IPTG was added to 0.5 mM. After four hours, cultures were centrifuged at 8200 x *g*, 4°C for 5 minutes, the supernatant was poured off, and cells were frozen at -80°C until purification. The day of purification, cells were thawed in Buffer A (50 mM Tris•Cl, pH 7.5, 150 mM NaCl, 5 mM MgCl_2_, 10% glycerol) supplemented with 20 mM imidazole, and lysed with an Avestin Emulsiflex C-5. Lysates were clarified via centrifugation at 30,000 x *g*, 4°C for 20 minutes, and the supernatant was incubated with Ni-NTA resin for 30 minutes at 4°C. After two washes with Buffer A containing 20 mM imidazole, proteins were eluted with Buffer A containing 500mM imidazole. The eluate was concentrated using a 10,000 Da MWCO Amicon filter, and further purified by gel filtration on a Sephacryl S-200 column in Buffer A.

### Purification of Sem1

Sem1 was expressed as 6His fusions in bacterial strain LOBSTR (DE3) co-transformed with pRARE2. Transformants were grown in LB and the appropriate antibiotics at 37°C, 250 rpm shaking until OD_600_ ≈ 1.0, at which point the temperature was reduced to 16°C and IPTG was added to 0.2 mM. After four hours, cultures were centrifuged at 8200 x *g*, 4°C for 5 minutes, the supernatant was poured off, and cells were frozen at -80°C until purification. The day of purification, cells were thawed in Sem1 Buffer (50 mM Tris•Cl, pH 7.5, 150 mM NaCl, 10% glycerol) supplemented with 2 mM PMSF and lysed by adding the cells dropwise to boiling Sem1 Buffer. The lysate was transferred to an ice water bath for 10 minutes before being centrifuged at 30,000 x *g* for 20 minutes. The supernatant was incubated with Ni-NTA resin for 30 minutes at room temperature. After two washes with Sem1 Buffer supplemented with 10 mM imidazole, proteins were eluted with Sem1 Buffer supplemented with 250 mM imidazole. The eluate was loaded onto a Mono Q 5/5 column equilibrated in Sem1 IX-A Buffer (50 mM Tris•Cl, pH 7.5), and proteins were eluted by a continuous gradient of Sem1 IX-B Buffer (50 mM Tris•Cl, pH 7.5, 1 M NaCl).

### Purification of proteasome substrates and Ubp6

All proteasome substrates and Ubp6 were expressed as 6His fusions in bacterial strain LOBSTR (DE3) co-transformed with pRARE2. Transformants were grown in 2 L of LB and the appropriate antibiotics at 37°C, 250 rpm shaking until OD_600_ ≈ 0.6, at which point the temperature was reduced to 16°C and IPTG was added to 0.5 mM. After overnight induction, cultures were centrifuged at 8200 x *g*, 16°C for 5 minutes, the supernatant was poured off, and cells were frozen at -80°C until purification. The day of purification, cells were thawed in NPI-10 (10 mM sodium phosphate, pH 8.0, 300 mM NaCl, 10 mM imidazole), and lysed with an Avestin Emulsiflex C-5. Lysates were clarified via centrifugation at 30,000 x *g*, 4°C for 20 minutes, and the supernatant was incubated with Ni-NTA resin for 30 minutes at 4°C. After two washes with NPI-10, the resin was poured into a disposable Bio-Rad Econo-column, washed with NPI-20 (10 mM sodium phosphate, pH 8.0, 300 mM NaCl, 20 mM imidazole), and eluted with NPI-500 (10 mM sodium phosphate, pH 8.0, 100 mM NaCl, 500 mM imidazole). Eluates were concentrated using 10,000 Da MWCO Amicon filters, and further purified by gel filtration on a Sephacryl S-200 column in Substrate Buffer (50 mM HEPES•OH, pH 7.5, 100 mM NaCl, 100 mM KCl, 5% glycerol). When present, fusion tags were cleaved either via an overnight incubation at 4°C with HRV-3C protease (1:20 w/w) or *S. cerevisiae* Ulp1 protease (1:1000 w/w) prior to gel filtration.

### Fluorophore labeling of proteasome substrates

Substrates were diluted to 100 µM in Substrate Buffer supplemented with 0.1 mM TCEP and incubated with a 5-fold molar excess of maleimide conjugate (sulfo-Cy3, sulfo- Cy5, or TAMRA as indicated) for 2 hours at room temperature. The reaction was quenched with 50 mM DTT for 30 minutes at room temperature. Excess dye was removed by desalting using either a PD-10 gravity column (Cytiva Life Sciences) or a Zeba desalting spin column (Thermo-Fisher Scientific).

### Purification of ubiquitin

Ubiquitin was expressed with an HRV3C-cleavable 6His tag at the N-terminus in bacterial strain LOBSTR (DE3) co-transformed with pRARE2LysS. Transformants were grown in 2xYT and the appropriate antibiotics at 37°C, 250 rpm shaking until OD_600_ ≈ 0.6, at which point IPTG was added to 0.4 mM. After 4 hours of induction at 37°C, cultures were centrifuged at 8200 x *g*, 16°C for 5 minutes, the supernatant was poured off, and cells were frozen at -80°C until purification. The day of purification, cells were thawed in NPI-10 and lysed with an Avestin Emulsiflex C-5. Lysates were clarified via centrifugation at 30,000 x *g*, 4°C for 20 minutes, and the supernatant was incubated with Ni-NTA resin for 30 minutes at 4°C. After two washes with NPI-10, the resin was poured into a disposable Bio-Rad Econo-column, washed with NPI-20, and eluted with NPI-500. HRV- 3C protease was added 1:20 (w/w) to the eluate, and it was dialyzed overnight at 4_o_C using 3500 MWCO dialysis tubing in 50 mM Tris•Cl, pH 8.0. The following morning, the dialysate was adjusted to pH 4.5 with acetic acid and purified by cation exchange on a HiTrap SP column equilibrated in Ubiquitin Cat-A buffer (50 mM ammonium acetate, pH 4.5). Proteins were eluted by a continuous gradient of Ubiquitin Cat-B Buffer (50 mM ammonium acetate, pH 4.5, 1 M NaCl). Pure fractions were combined and dialyzed as above into Ubiquitin Storage Buffer (20 mM Tris•Cl, pH 7.5, 0.5 mM EDTA).

### Purification of ubiquitination machinery

Enzymes for *in vitro* ubiquitination (mouse E1, yeast Ubc1, and yeast Rsp5) were expressed as 6His or 12His fusions in bacterial strain LOBSTR (DE3) co-transformed with pRARE2. Transformants were grown in LB and the appropriate antibiotics at 37°C, 250 rpm until OD_600_ ≈ 0.6, at which point the temperature was reduced to 16°C and IPTG was added to 0.5 mM. After overnight induction, cultures were centrifuged at 8200 x *g*, 16°C for 5 minutes, the supernatant was poured off, and cells were frozen at -80°C until purification. The day of purification, cells were thawed in NPI-10 and lysed with an Avestin Emulsiflex C-5. Lysates were clarified via centrifugation at 30,000 x *g*, 4°C for 20 minutes, and the supernatant was incubated with Ni-NTA resin for 30 minutes at 4°C. After two washes with NPI-10, the resin was poured into a disposable Bio-Rad Econo-column, washed with NPI-20, and eluted with NPI-500. Eluates were concentrated using 10,000 Da MWCO Amicon filters, and further purified by gel filtration on a Sephacryl S-200 column in Ub/DUB Buffer (50 mM Tris•Cl, pH 7.5, 100 mM NaCl).

### Substrate ubiquitination

Ubiquitination of substrates was carried out with 20-50 µM substrate incubated with 5 µM mouse E1, 5 µM Ubc1, 2.5 µM Rsp5, 0.5-1 mM ubiquitin, 15 mM ATP, and 600 µM DTT in Ubiquitination Reaction Buffer (50 mM Tris•Cl, pH 8.0, 5 mM MgCl_2_) at 25°C for at least 2 hours. Successful substrate ubiquitination was confirmed via SDS-PAGE prior to use in experiments.

### Substrate degradation assays

Degradation assays using GFP substrates were performed at 30°C in 26S Buffer (50 mM Tris•Cl, pH 7.5, 50 mM NaCl, 50 mM KCl, 5 mM MgCl_2_, 1 mM EDTA, 10% glycerol, 1 mM DTT) with an ATP-regenerating system (60 µg/mL creatine kinase, 16 mM creatine phosphate, 5 mM ATP) and in the presence of 0.4 mg/mL BSA. Proteasomes were reconstituted from 50 nM CP, 100 nM base, 100 nM Rpn10, 400 nM WT or sem1Δ lid, and 600 nM Sem1 where indicated. Excess lid did not affect observed degradation rates. Degradation assays were initiated by addition of the indicated concentration of GFP substrate, and degradation was monitored by the loss of GFP fluorescence (ex 479 nm, em 520 nm) on a BioTek Synergy H1MF multi-mode reader. Linear regression was performed on the fluorescence traces to determine the initial reaction velocities. For Michaelis-Menten analyses, the initial rates were plotted against substrate concentration and fitted to the Michaelis-Menten equation using Prism 8 (Graphpad Software, Inc.) to determine *k*_cat_ and *K*_M_.

### Substrate tail engagement assays

Assays were performed essentially as described by (Bard, et al., 2019). Substrate tail insertion was measured by monitoring Förster resonance energy transfer (FRET) between Cy3-labeled base and Cy5-labeled ubiquitinated substrate using a KinTek AutoSF-120 stopped-flow system. Single-turnover conditions were enforced using *ortho*- phenanthroline, a zinc chelator that inactivates Rpn11, thus stalling the proteasome at the site of ubiquitin linkage. Reactions were performed by rapid mixing 2x concentrations of reconstituted proteasomes with substrate. Final concentrations were 100 nM Rpt1-^I191AzF-Cy3^ base, 400 nM CP, 600 nM WT or sem1Δ lid, 750 nM Rpn10, 3 mM *o*-PA, and 3 µM ubiquitinated Cy5-substrate. Samples were excited at a wavelength of 532 nm, and FRET was monitored by simultaneous detection of Cy3 (595 nm, 50 nm bandpass filter) and Cy5 (690 nm, 50 nm bandpass filter) emission. Kinetics were determined by fitting the change in the intensity of the Cy3 signal to a one-phase association, which was confirmed by analysis of residuals to yield the best fit.

### Ubp6 Ub-AMC cleavage-activity assays

Ubp6 deubiquitination activity was measured by mixing reconstituted proteasomes under base-limiting conditions (200 nM CP, 100 nM base, 400 nM Rpn10, 400 nM WT or sem1Δ lid, and 2 µM Sem1 where indicated in 26S Buffer) with 40 nM Ubp6, 0.1 mg/mL BSA, and 5 mM ATP or ATPγS. After 15 minutes at 30°C, the fluorogenic Ub-AMC substrate (Enzo Life Sciences) was added to a final concentration of 1 µM. Cleavage was measured by monitoring the change in fluorescence intensity (ex 360 nm, em 460 nm) on a BioTek Synergy H1MF multi-mode reader. Relative rates were determined from the initial slopes of fluorescence versus time, and the fold change compared to WT 26S in ATP was plotted.

### Deubiquitination assays

Assays were performed essentially as described (Bard, et al., 2019). Deubiquitination rates were determined by monitoring FRET between a Cy5-labeled titin-I27^V15P^ substrate that has been modified with Cy3-labeled ubiquitin using a KinTek AutoSF-120 stopped-flow system as described above. Final concentrations were 400 nM CP, 100 nM base, 500 nM WT or sem1Δ lid, 750 nM Rpn10, 750 nM Sem1 (where noted), and 20 nM ubiquitinated substrate. Kinetics were determined by fitting the change in the intensity of the Cy5 signal to a one-phase decay, which was confirmed by analysis of residuals to yield the best fit.

### Peptidase stimulation assays

Peptidase stimulation was measured by cleavage of the fluorogenic substrate suc-LLVY-AMC. Proteasomes were reconstituted from 10 nM CP, 80 nM base, 80 nM Rpn10, 300 nM WT or sem1Δ lid, and 1.5 µM Sem1 where indicated in 26S Buffer supplemented with 5 mM ATP and 0.1 mg/mL BSA. After 15 minutes at 30°C, suc-LLVY-AMC was added to a final concentration of 50 µM. Cleavage was measured by monitoring the change in fluorescence intensity (ex 360 nm, em 460 nm) on a BioTek Synergy H1MF multi-mode reader. Relative rates were determined from the initial slopes of fluorescence versus time, and plotted as percent of CP alone.

### ATPase activity assays

The ATP hydrolysis rate of reconstituted proteasomes was measured using a NADH-coupled kinetic assay (Nørby, 1988). Proteasomes were reconstituted under base-limiting conditions (400 nM CP, 200 nM base, 800 nM Rpn10, 800 nM WT or sem1Δlid, 4 µM Sem1 where indicated) in 26S buffer and supplied with 1 µM Ub_4_-GFP-Tail substrate where indicated. ATPase mix was added (final concentrations: 1.5 U/mL pyruvate kinase, 1.5 U/mL lactate dehydrogenase, 3.75 mM phospho(enol)pyruvic acid, 1 mM NADH, and 2.5 mM ATP), and the change in absorbance at 340 nm was monitored on a BioTek Synergy H1MF multi-mode reader.

### Rpn11 restart assays

Assays were performed similarly to those described (Greene, et al., 2019). Proteasomes (100 nM CP, 200 nM base, 200 nM Rpn10, 800 nM lid, and 1.2 µM Sem1 where indicated) were reconstituted in 26S buffer with 5 mM ATP and 1 mM DTT and allowed to assemble for 15 minutes at 30°C. Proteasomes were then stalled with 3 mM *ortho*-phenanthroline, followed by addition of 20 nM substrate (GFP-polyUb-Tail) and incubation at room temperature for 3 minutes. Stalled proteasomes were restarted by adding ZnSO_4_ to a final concentration of 1 mM. Samples were immediately transferred to a BioTek Synergy H1MF multi-mode reader to monitor loss of GFP fluorescence. Fluorescence intensity versus time was plotted and fit to a first order exponential to derive unfolding rates.

### Unfolding ability assays

Assays were performed similarly to those described (Cresti, et al., 2021; Hurley & Kraut, 2021). Neh2Dual-BarnaseΔK(L89G)-TAMRA-DHFRδ5KΔC was ubiquitinated using the following conditions: 1.5 µM substrate, 1.5 µM E1, 3 µM Ubc1, 3 µM Rsp5, 1.5 mg/mL ubiquitin, 5 mM ATP, 1 µM DTT, and 5 µM methotrexate in ubiquitin reaction buffer for 2 h at 25°C. Proteasomes were reconstituted from 100 nM CP, 200 nM base, 200 nM Rpn10, 800 nM WT or sem1Δ lid in the presence of 1 µM methotrexate and unfolding assay buffer (50 mM Tris•Cl, pH 8.0, 5 mM MgCl_2_, 5% glycerol). Ubiquitinated substrate was added to a concentration of 40 nM. Aliquots were taken at the indicated timepoints and placed into SDS-PAGE loading buffer to quench the reaction. Samples were loaded onto a 9.25% tricine gel, and fluorescence was visualized using a Bio-Rad ChemiDoc MP. Bands corresponding to full-length substrate and DHFR-containing fragments were quantified and fit to single exponential equations to determine the amplitudes of degradation and fragment formation. The amplitudes were used to calculate unfolding ability (U) using the following equation:

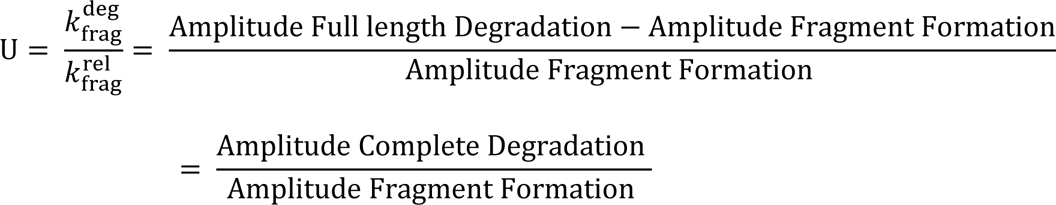

### Conformation-selective engineered disulfide crosslinking

Crosslinking was performed essentially as described previously (Eisele, et al., 2018; Reed & Tomko, 2019). Yeast expressing proteins with the desired cysteine substitutions were grown to mid-log phase, 20 OD_600_ equivalents were harvested, and the cell were converted to spheroplasts. Spheroplasts were lysed in 150 µL of ice-cold lysis buffer (50 mM HEPES•OH, pH 7.5, 150 mM NaCl, 5 mM MgCl_2_) containing 2 mM of nucleotide (ATP or AMP-PNP). The cells were lysed by vortexing at top speed three times for 30 seconds with 1-minute rests on ice in between. The lysates were centrifuged at 21,000 x *g*, 4°C for 10 minutes to clear cell debris. The protein content of the supernatant was normalized with lysis buffer containing the appropriate nucleotide. Crosslinking was initiated with 250 µM CuCl_2_ at 25°C. After 10 minutes, crosslinking was quenched with 10 mM *N*-ethylmaleimide and 10 mM EDTA. To reduce the engineered disulfides prior to SDS-PAGE analysis, DTT was added to a final concentration of 20 mM, and samples were incubated at room temperature for 10 minutes. Samples were then boiled in non-reducing Laemmli buffer, loaded onto 10% SDS-PAGE gels, and separated by electrophoresis before immunoblotting with antibodies against V5 or G6PD.

### Quantification and statistical analysis

All experiments were performed at least twice, with most experiments being repeated three or more times. Statistical analysis was carried out using Graph Pad Prism 8.0 software using one- or two-way ANOVA where appropriate with Tukey’s or Sidak’s tests for multiple comparisons. Exact values of statistical significance (*p*) and *N* for each experiment can be found in the figure legends.

## ACKNOWLEDGEMENTS

The authors thank Mark Hochstrasser (Yale University) for anti-Sem1 antibodies and the Tomko lab for helpful feedback on the manuscript. This work was supported by NIH grant 5R01GM118600 to R.J.T.

## AUTHOR CONTRIBUTIONS

R.G.R: Conceptulization, formal analysis, investigation, methodology, resources, validation, visualization, writing—original draft. G.W.J: investigation, resources, writing— review and editing. R.J.T: Conceptualization, formal analysis, funding acquisition, investigation, methodology, project administration, resources, supervision, writing— review and editing.

---

**Figure 1 – Figure Supplement 1.**
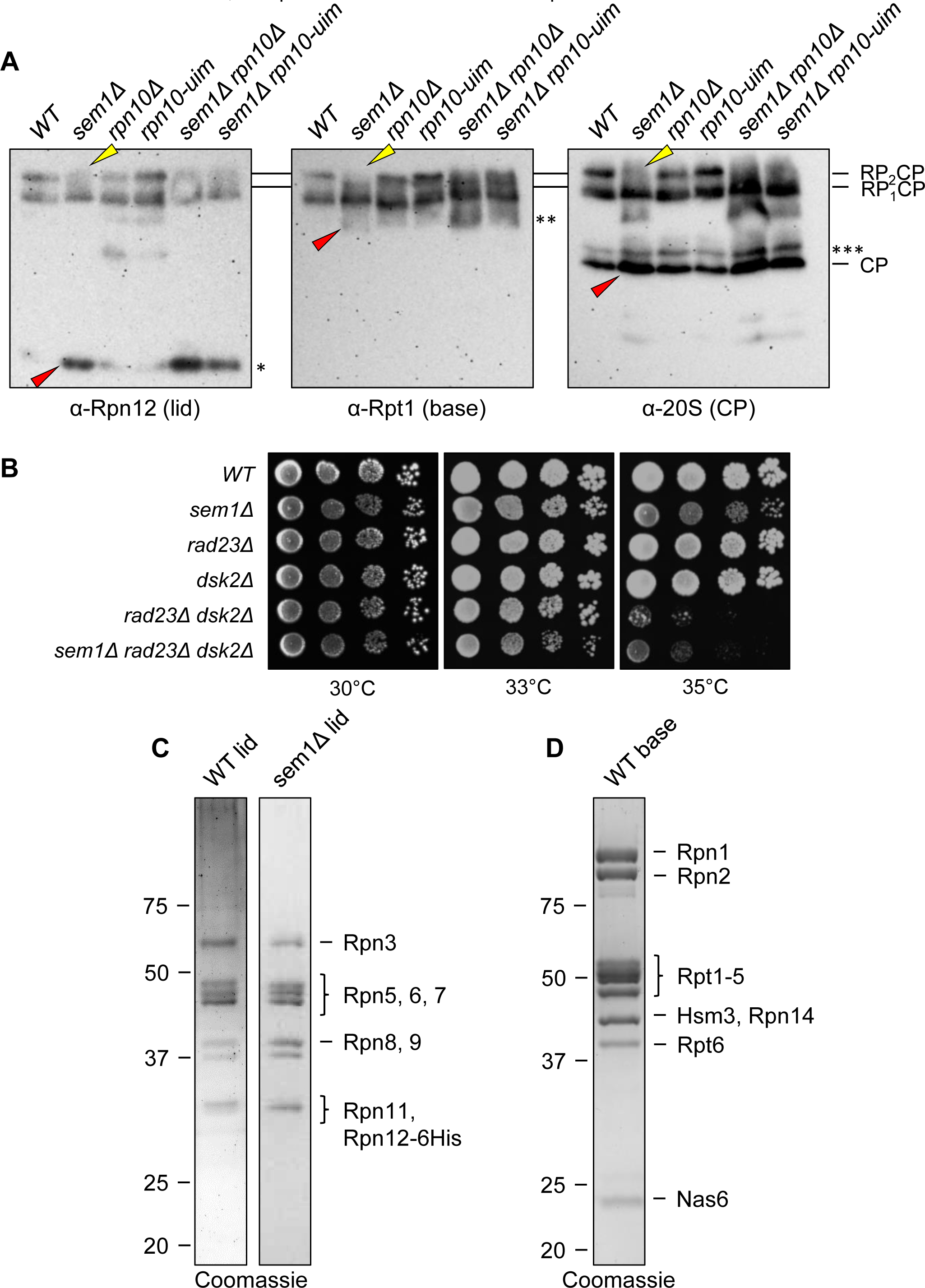
Structural defects of *sem1Δ rpn10Δ* proteasomes and purified lid and base sub-complexes. (A) Extracts from the indicated yeast strains were separated by native PAGE and immunoblotted with antibodies against Rpn12, Rpt1, or the CP. Yellow arrowheads indicate decrease of doubly capped proteasomes (RP_2_CP) in *sem1Δ* extracts compared to *WT*. Red arrowheads indicate accumulation of free lid subunit Rpn12, base subcomplex, and CP subcomplex in *sem1Δ* extracts compared to WT. *, free Rpn12. **, free base. ***, intermediate complex. (B) Deletion of *SEM1* fails to yield synthetic growth phenotypes when combined with proteasomal substrate shuttle factor deletions. Equal numbers of cells with the indicated genotypes were incubated on YPD medium for 3 days at the temperatures shown. (C) WT or sem1Δ purified recombinant lids and (**D**) WT recombinant base were separated by SDS-PAGE and stained with Coomassie.

**Figure 3 – Figure Supplement 1.**
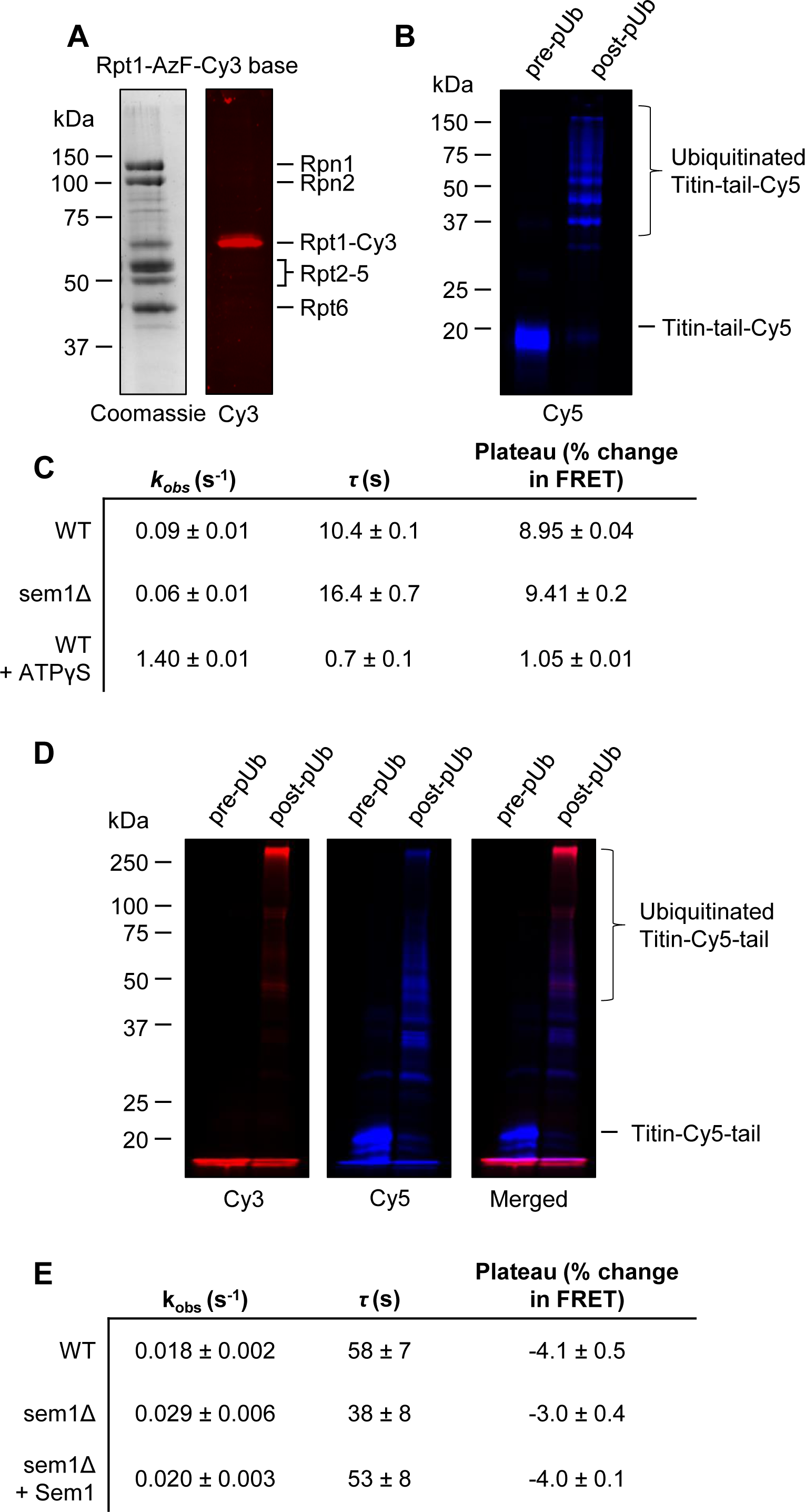
Substrate tail engagement and deubiquitination assay components and results. (A) Site-specific labeling of Rpt1-AzF-Cy3 base. The position of each subunit of the base subcomplex is indicated. (B) Polyubiquitination of Cy5-labeled titin substrate. Cy5-labeled Titin-Tail was ubiquitinated using recombinant Ub, mmE1, Ubc1, and Rsp5 before analysis by SDS-PAGE. (C) Derived values for *k*_obs_, τ, and plateau from the data fit shown in Figure 3C. Error represents SEM. (D) Polyubiquitination of Cy5-labeled titin substrate with Cy3-labeled ubiquitin. (E) Derived values for *k*_obs_, τ, and plateau from data fit in Figure 3F. Error represents SEM.

**Figure 5 – Figure Supplement 1.**
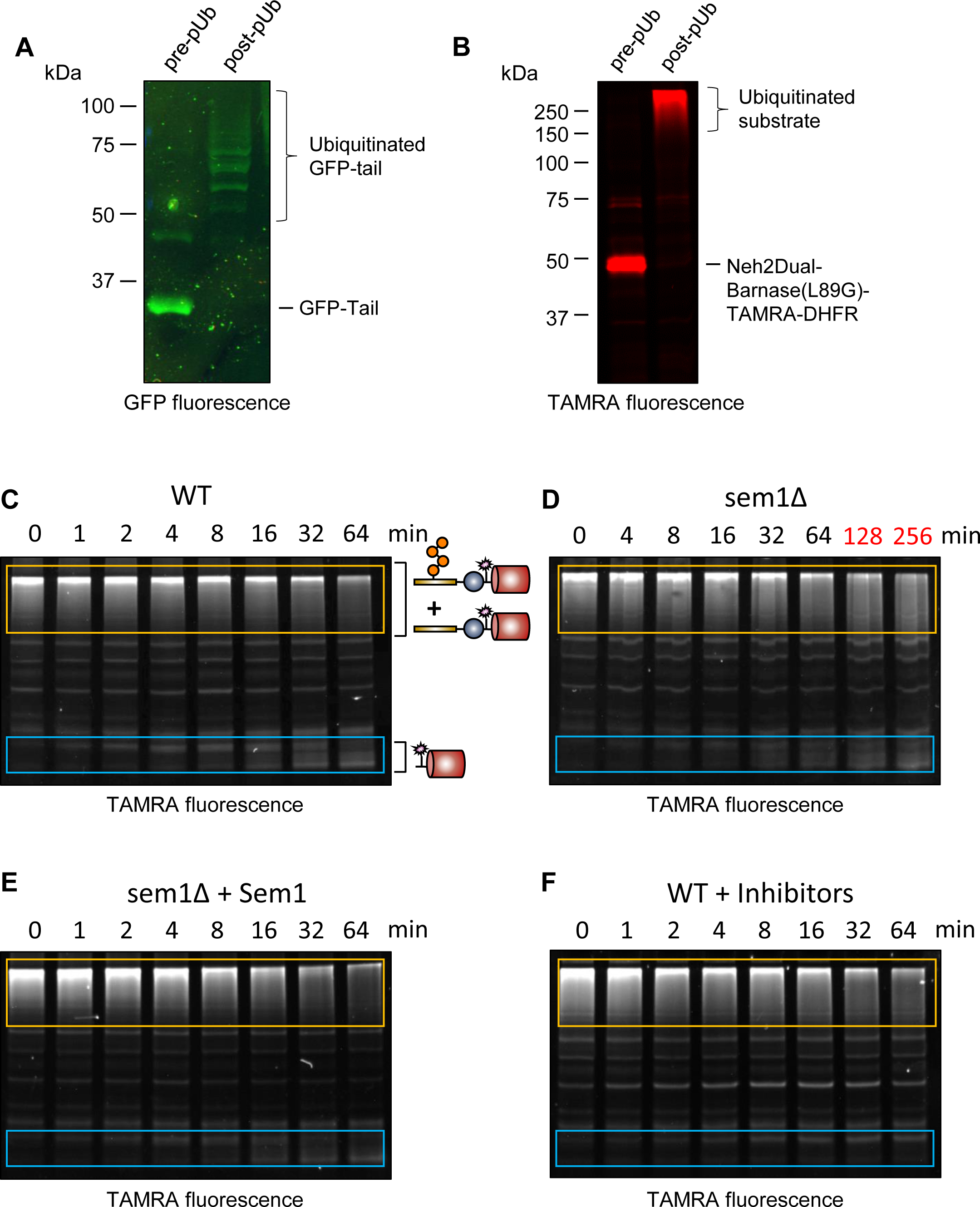
Proteasomes reconstituted without Sem1 harbor an unfolding defect. (A) Polyubiquitination of GFP-Tail substrate. (B) Polyubiquitination of Neh2Dual-Barnase(L89G)-TAMRA-DHFR. (**C-F**) Degradation of polyubiquitinated Neh2Dual-Barnase(L89G)-TAMRA-DHFR by WT (**C**), sem1Δ (**D**), sem1Δ + Sem1 (**E**), or WT proteasomes + inhibitors MG132, bortezomib, and *o*PA (**F**). Yellow boxes (top) surround the region of the lane containing full-length or deubiquitinated substrate. Blue boxes (bottom) surround the region of the lane containing the DHFR fragment. The quantifications of these bands are shown as percentages of the starting full-length + non-ubiquitinated substrate in Figure 5D.

**Figure 6 – Figure Supplement 1.**
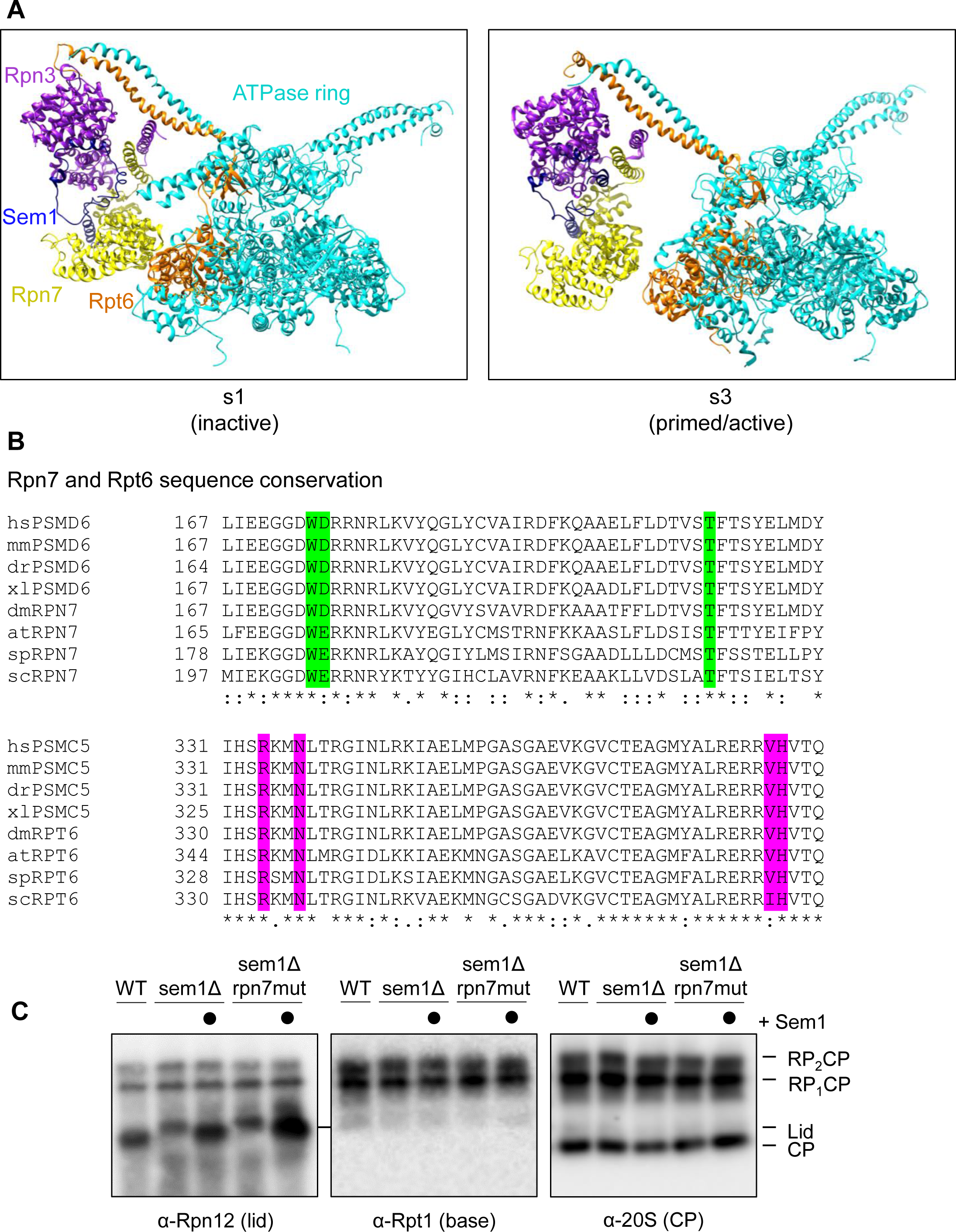
Sem1 does not make direct contact with the ATPase ring. (A) The proteasome ATPase ring and lid subunits Rpn3, Rpn7, and Sem1 in the inactive/s1 state (PDB: 6FVT) versus the primed/active/s3-like states (PDB: 6FVV). Several proteasome subunits are omitted for clarity. (B) Multiple species alignment of Rpn7 and Rpt6 regions contributing to the contacts seen in Figure 6A. hs, *Homo sapiens*; mm, *Mus musculus*; dr, *Danio rerio*; xl, *Xenopus laevis*; dm, *Drosophila melanogaster*; at, *Arabidopsis thaliana*; sp*, Schizosaccharomyces pombe*; sc, *Saccharomyces cerevisiae*.

**Figure 7 – Figure Supplement 1.**
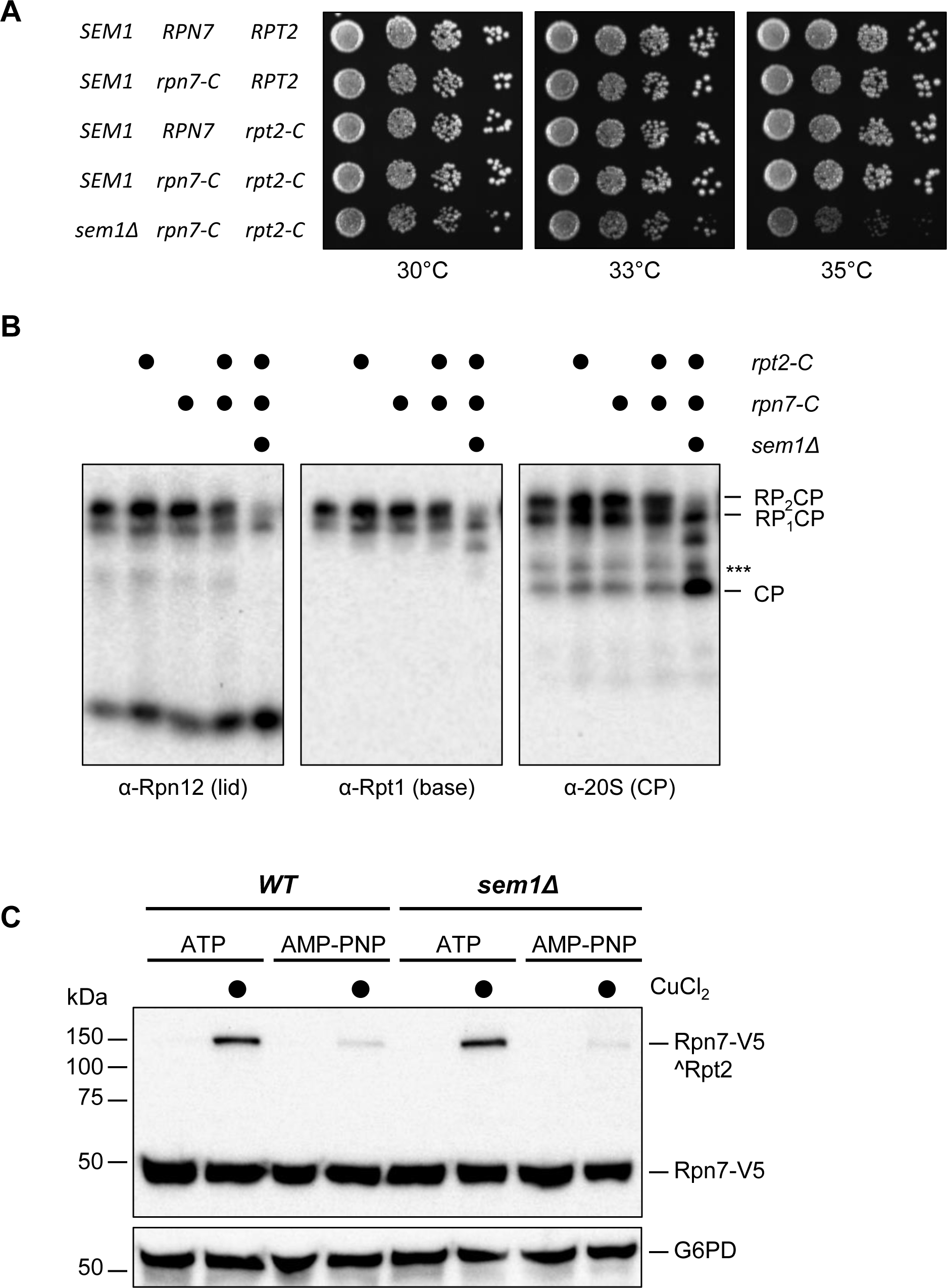
Crosslinking of Rpn7 and Rpt2 in a *sem1Δ* strain. (A) Equal numbers of cells with the indicated genotypes were incubated on YPD medium for 2 days at the temperatures shown. (B) Extracts from the indicated yeast strains were separated by native PAGE and immunoblotted with antibodies against Rpn12, Rpt1, or the CP. ***, intermediate complex. (C) Representative immunoblot of Rpn7/Rpt2 crosslinking of *WT* and *sem1Δ* WCEs quantified in Figure 7A.

**Figure 7 – Figure Supplement 2.**
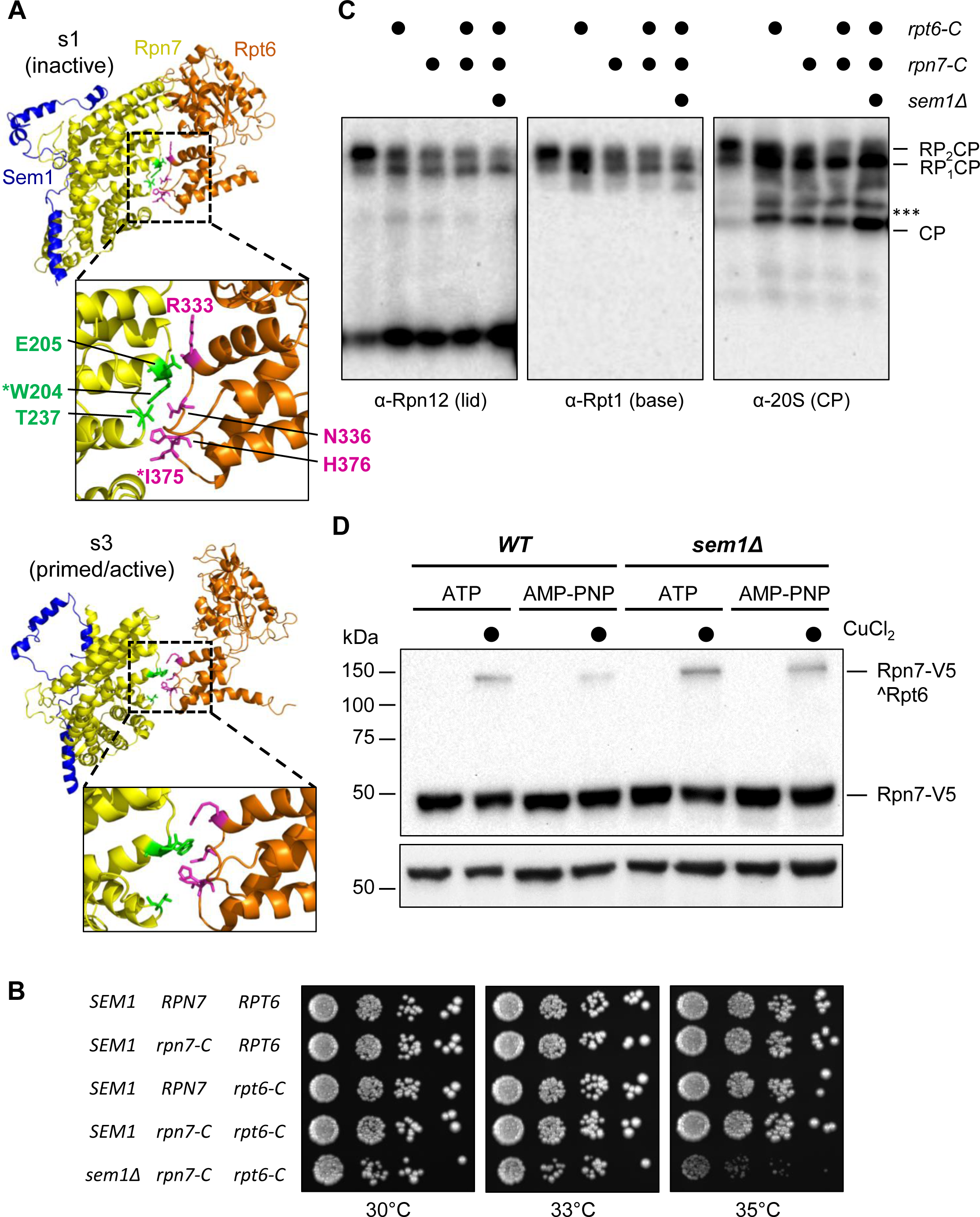
Crosslinking of Rpn7 and Rpt6 in a *sem1Δ* strain. (A) A close view of the residues making contact between Rpn7 and Rpt6 in the inactive s1 state (PDB: 6FVT) and the active s3 state (PDB: 6FVV). Residues Rpn7 W204 and Rpt6 I375 (marked with asterisks) were mutated to cysteines for disulfide crosslinking experiments. (B) Equal numbers of cells with the indicated genotypes were incubated on YPD medium for 3 days at the temperatures shown. (C) Extracts from the indicated yeast strains were separated by native PAGE and immunoblotted with antibodies against Rpn12, Rpt1, or the CP. ***, intermediate complex. Representative immunoblot of Rpn7-Rpt6 crosslinking of *WT* and *sem1Δ* WCEs quantified in Figure 7B.

